# Sex-linked dilution colour in the European domestic goose confirmed to be a 1-bp deletion in the *Melan-A* gene

**DOI:** 10.64898/2026.01.19.700295

**Authors:** Suvi Olli, Vikke Ahola, Marja E. Heikkinen, Johanna Honka

## Abstract

Plumage colour in domestic geese is an important economic trait and a selection target since the early days of domestication. In European domestic geese of greylag goose (*Anser anser*) origin, white plumage colour is known to be caused by two independent loci, one causing white spotting and one causing sex-linked dilution, together producing white plumage. Strong candidate mutations have been identified upstream of the *EDNRB2/LOC106047519* gene (*endothelin receptor B-like*) and within the sex-linked *MLANA* gene (*melan-A*). To confirm these candidate mutations, we genotyped differently coloured European domestic goose breeds, wild greylag geese, Chinese domestic geese (derived from swan goose *A. cygnoid*) and European and Chinese domestic geese crossbreeds. One base pair deletion in the *MLANA* gene (NW_013185876.1: g.950,868 C > -) was confirmed to cause sex-linked dilution, and thus autosexing (almost white gander and goose diluted grey). However, mutation upstream of *EDNRB2/LOC106047519* (NW_013185915.1: g. 775,151 G > T) was not the causative mutation for saddleback pattern but strongly linked to it in European domestic geese. We sequenced the *EDNRB2* gene and coding sequence of a neighbouring *VAMP7* gene (*vesicle-associated membrane protein 7*) but found no genetic variaion linked to colour. Additionally, we sequenced the coding sequence of *TYRP1* (*tyrosinase related protein 1*), a candidate gene for buff colouration, but no variation linked to colour was found. Further, we genotyped a 14-bp insertion in exon 3 of the *EDNRB2* gene, known to be causative of the white phenotype in the Chinese domestic goose, and identified it in one European domestic goose individual.

## Introduction

White plumage in domestic geese has been attractive to humans since the beginning of the domestication of these species, the European domestic goose deriving from the greylag goose (*Anser anser*) and the Chinese domestic goose from the swan goose (*A. cygnoid*). The greylag goose has been speculated to be domesticated around 3000 BCE (Before Common Era; Albarella 2005) or about 5500 BCE based on genomic data of modern breeds, while the genomic estimate for the Chinese domestic goose was only 1500 BCE (Wen et al. 2023). The first archaeological evidence of domestic geese derives from 5000 BCE in China, but the species is unknown as both progenitor species occur in the area (Eda et al. 2022). The first evidence of white geese dates to the Egyptian New Kingdom times (1552–1151 BCE) from tomb wall art depicting flocks of variedly coloured geese, including white, grey, and geese with grey patches on the front or back of their head resembling modern saddleback/pied/mottled colouring (Fig. 1a and 1b; Bailleul-LeSuer 2012, Bailleul-LeSuer 2016). A pair of differently coloured parent geese followed by goslings is further proof of domestication (Fig. 1c; Bailleul-LeSuer 2012). As leucistic geese are rare in nature, pure white or mottled geese are a definitive sign of domestication, as the attractive colouration was selected by humans. Besides ornamental appearance, white geese also have faster growth rates, provide more attractive carcasses (lack of dark feather quills) and white down for stuffing, e.g. pillows and mattresses, and allegedly tastier meat (Kear 1990). However, the grey type starts to lay earlier and produces more eggs (Kear 1990).

**Fig. 1.**
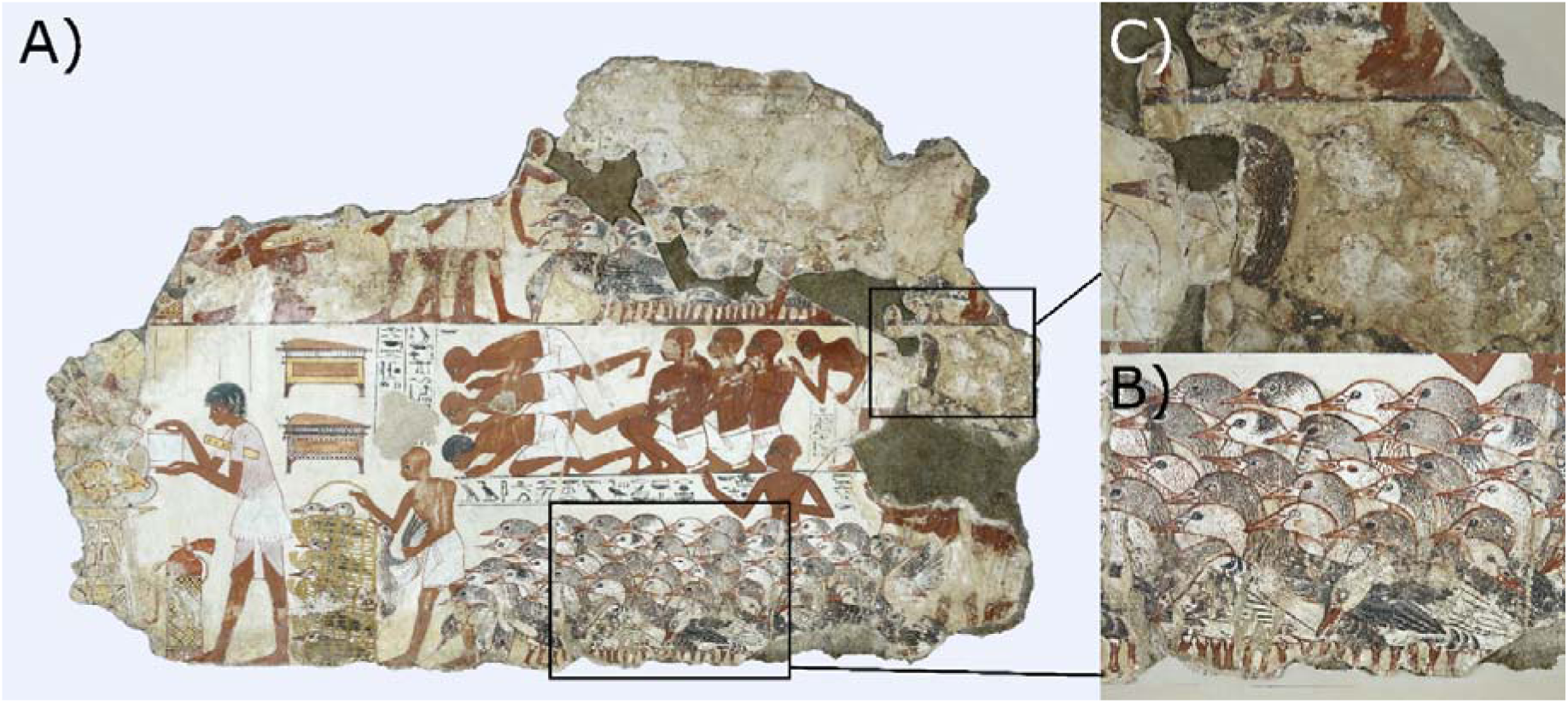
Flocks of heterogeneously coloured geese depicted in Egyptian 18^th^ Dynasty (1350 BCE) wall painting from the tomb-chapel of Nebanum. A) The whole piece of the wall painting, B) detail of the heterogeneously coloured goose flock and C) detail of a pair of differently coloured parent geese with their goslings. Photo courtesies: The Trustees of the British Museum (CC BY-NC-SA 4.0).

The European domestic goose exhibits several plumage colourations: wild type grey, white, saddleback also known as pied, piebald or mottled (referred as saddleback from hereon), autosexing (almost white male and light grey or saddleback female), buff (light brownish/beige colour) and blue (bluish/silvery grey) (Fig. 2). Saddleback pattern is caused by white spotting, and such geese are white except for grey (or buff or blue, depending on underlying colour) feathers on the back, head and thigh coverts (Fig. 2C). The extent of white and grey can vary (Fig. 2F) and is possibly determined by another locus. Geese heterozygous for the white spotting display a white chin spot, a white breast patch and/or white flight feathers (Fig. 2D), though the expression is highly variable. In autosexing breeds the sexes can be conveniently sorted by their plumage colour; gander being almost white with some grey feathers on the rump, wings and tail and female either diluted grey often having white ‘spectacles’ (Pilgrim, Cotton Patch and Settler goose), or light grey saddleback (West of England, Shetland, Normandy and Bavent goose) (Fig. 2E-2F). Even white European domestic goose goslings are autosexing as female goslings have a darker grey saddleback plumage pattern, while in male goslings the saddleback pattern is of lighter grey colour and usually does not reach the head (Fig. 3A). However, it has been noted that Embden goose goslings autosex more reliably than Czech goose goslings, as in Embden goslings the female fluff is of darker shade (Ashton & Ashton 2023). Small-scale breeding experiments with Czech geese have raised the possibility that selection of other modifying factors makes this breed “whiter” than the Embden breed (Ashton & Ashton 2023). As white breed goslings develop, both sexes turn white. In grey breeds, the goslings have grey fluff (Fig. 3B), the colour being distributed as in buff breeds (Fig. 3C). In autosexing breeds of the type ‘almost white male and light grey female’, the goslings also autosex with females having darker fluff than males (Fig. 3D). Goslings of the autosexing breeds of the type ‘almost white male and saddleback female’ show saddleback pattern, but the shades of grey are darker than in white breeds (Fig. 3E). In saddleback breeds, the saddleback pattern is visible from birth to adult.

**Fig. 2.**
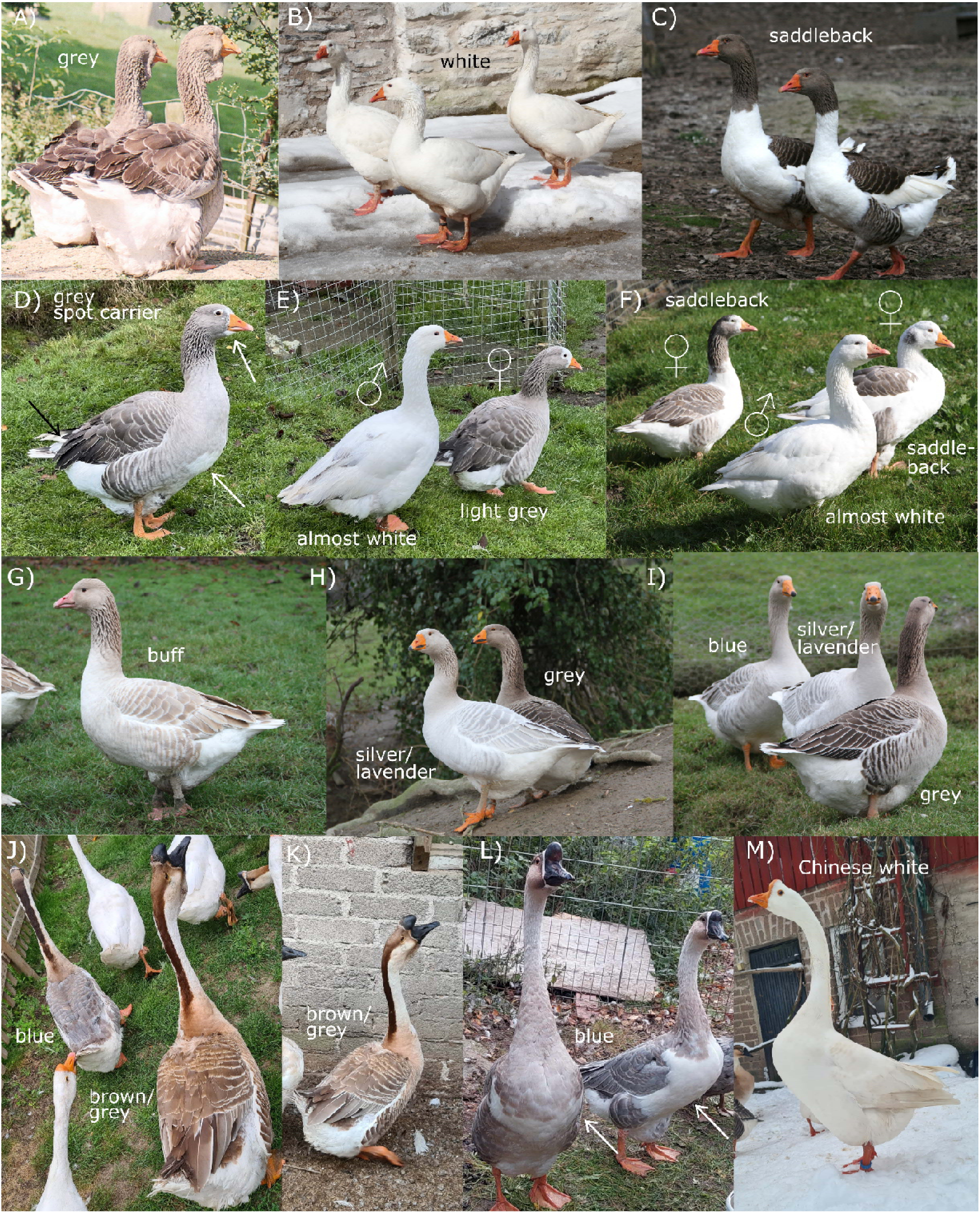
Phenotypes of the European domestic geese (*Anser anser* -derived) and Chinese domestic geese (*A. cygnoid* -derived). It should be noted that light exposure (bright daylight - overcast) varies between the photos. A) Grey wild type European domestic geese of Toulouse breed, B) white European domestic geese of Embden breed, C) saddleback European domestic geese of Pomeranian breed, D) grey spot carrier (heterozygous for spotting) from a cross between Shetland gander and Brecon Buff goose, note a white breast patch, a white chin and white flight feathers, E) autosexing Pilgrim goose breed with almost white gander and diluted grey female, often having white feathers in the front of the head, F) autosexing Shetland goose breed with almost white gander and saddleback females, G) buff European domestic goose of Brecon Buff breed, H) silver/lavender (homozygous for blue allele) Steinbacher goose (foreground) and grey Steinbacher in the background, Steinbacher is a cross between European domestic and Chinese domestic goose, I) blue Steinbacher (further back), silver/lavender Steinbacher (middle) and grey Steinbacher (foreground), J) blue Chinese domestic goose in the left and brown a.k.a. grey Chinese domestic goose in the right showing the side-by-side colour comparison, K) brown a.k.a. grey Chinese domestic goose, L) two blue Chinese domestic geese carrying one white allele (heterozygous for white colour), showing variable sizes of white breast patches, and M) white Chinese domestic goose. Photos A-I: Chris and Mike Ashton, photos J, K and M: Kirsi Mäki, photo L: Heini Koskinen.

**Fig. 3.**
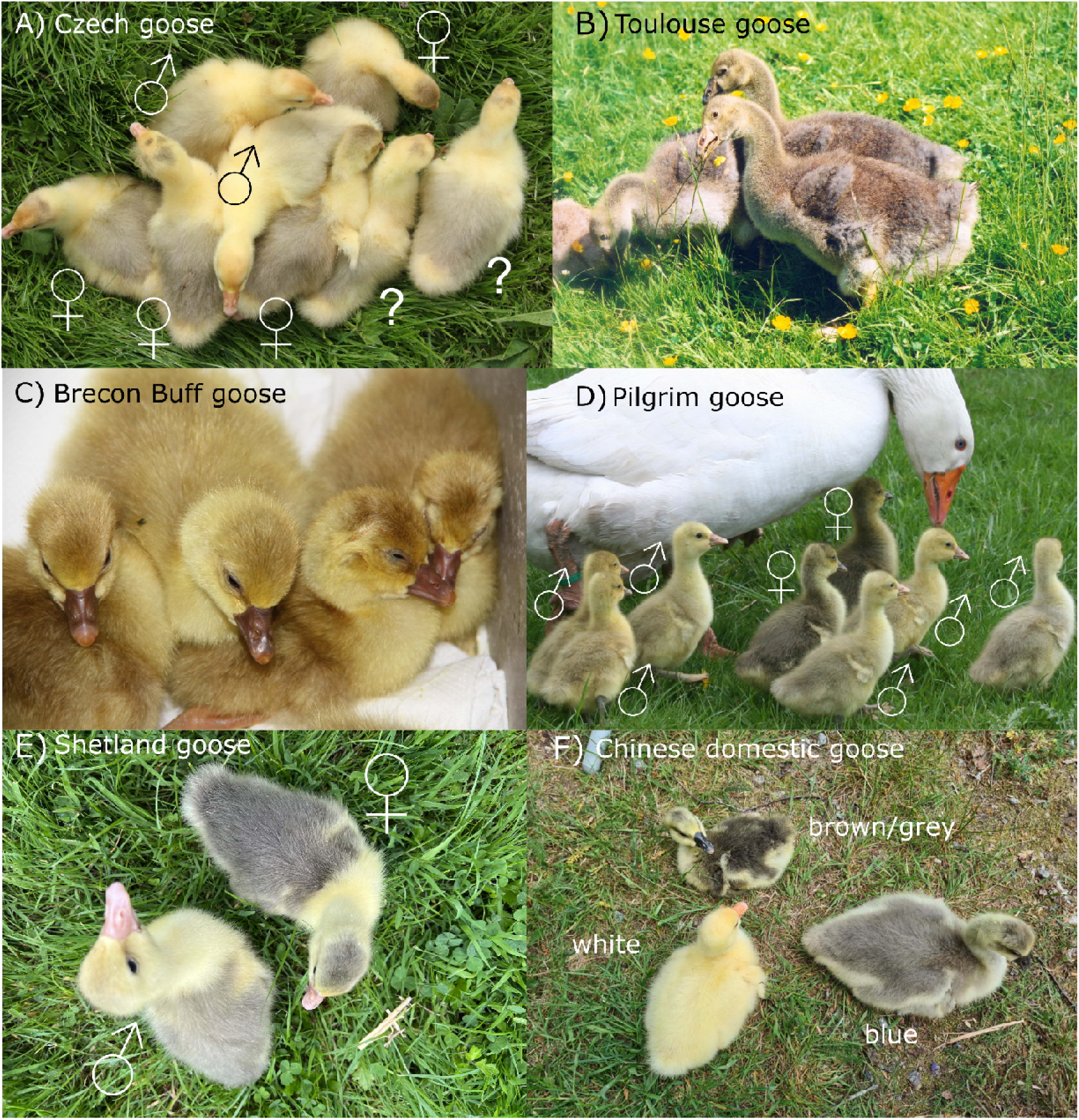
European domestic geese (*Anser anser* -derived) and Chinese domestic geese (*A. cygnoid* - derived) gosling plumages. White European domestic goose breeds are autosexing based on gosling plumage, females showing a darker grey saddleback pattern in contrast to a lighter shade of grey in males. In autosexing breeds, the sex of the individual can be differentiated both in the fluff and as adults. A) White breed (Czech goose) goslings, four females, two males and two goslings with unknown sex, as sexing in fluff is not 100% certain. B) Grey breed goslings (Toulouse goose) show uniform grey fluff in the back. C) Buff breed (Brecon buff) goslings also show uniform colour in the back. D) Autosexing breed in which the male is almost white and the female light grey (Pilgrim goose), goslings show darker shades of grey than white goose goslings, with darker grey females and lighter grey males. E) Autosexing breed in which the male is almost white and the female saddleback (Shetland goose), show saddleback pattern as goslings, with darker female and lighter grey male. F) White Chinese domestic goose goslings are completely yellow, brown/grey Chinese domestic goose goslings have dark grey fluff in the back and in blue gosling the grey is of a lighter colour. Photos A-E: Chris and Mike Ashton, photo F: Kirsi Mäki.

The inheritance patterns for the white phenotypes in geese were laid by Jerome (1953) by crossing white Embden geese with white Chinese domestic goose ganders, and based on his crosses, white European domestic goose has an allele for sex-linked dilution and white spotting, together producing white plumage. Geese with only spotting are saddlebacks, and geese with only sex-linked dilution have males diluted to almost white and females to light grey (Ashton 2021). In birds, female is a heterogametic sex with ZW sex chromosomes, while male has ZZ sex chromosomes and thus males are “double diluted” (homozygous) compared to hemizygous females. Underlying spotting is visible in white European domestic goose goslings as saddleback patterned fluff (Fig. 3A), which in adults is diluted to white due to the Z chromosomal dilution. As autosexing geese with ‘almost white male and saddleback female’ do not fit the expectations, these breeds are suggested to carry modified sex-linked dilution, which does not whiten areas over the female grey saddleback pattern (Ashton 2021). White Chinese is a colourless breed carrying an allele for non-colour (Jerome 1953) and lacks melanin pigment (producing white plumage). White Chinese domestic goose goslings have yellow fluff, while the fluff is darker in wild type brown aka grey individuals and lighter grey in blue individuals (Fig. 3F).

The genetic basis of white plumage has been studied using GWAS (genome-wide association studies) and selective sweep analysis. Strong candidate mutation for white spotting (saddleback) occurs upstream of the *EDNRB2* (*endothelin receptor B-like*) gene (NW_013185915.1: g. 775,151 G > T) (Yang et al. 2022). In the current reference genome (NC_089885.1), this gene is named as *LOC106047519* (*endothelin receptor type B-like*), but for simplicity, we refer to this as *EDNRB2* throughout this publication. In grey geese, allele T was fixed while allele G was fixed in white geese, possibly causing time-dependent downregulation of melanogenesis-related genes (Yang et al. 2022), supported also by findings of Liu et al. (2024). Melanoblasts (precursor cells of melanin-containing and producing melanocytes) originate from the embryonic neural crest from specific locations along the back and head, and these cells need to reach their allocated position in a certain developmental time window, and failing to do so, regions furthest away from the melanoblast origin, such as legs, belly, and forehead are often unpigmented (Cieslak et al. 2011). *EDNRB2* mutations have been identified to cause spotted/piebald or white phenotypes in several avian species (Miwa et al. 2007, Kinoshita et al. 2014, Xi et al. 2021, Nannan et al. 2024, Wang et al. 2025).

Downstream of the *EDNRB2* gene (9,967 bp between the genes) lies another possible gene contributing to melanosome transportation: *VAMP7* (*vesicle-associated membrane protein 7*). VAMP7 is a key protein in melanosomal transport, pigmentation and biogenesis as it regulates melanosome maturation together with STX13 (syntaxin13) by controlling endosomal cargo transport to the melanosomes (Jani et al. 2015). In the Chinese domestic goose, *VAMP7* has been identified as a candidate gene for melanin pigmentation (Zhao et al. 2025). We also studied this gene due to its proximity to the *EDNRB2* gene and role in melanosomal transportation.

A strong candidate mutation for sex-linked dilution is a 1-bp deletion (NW_013185876.1: g.950,868 C > -) in exon 4 of the Z chromosomal *MLANA* gene (*melan-A*), causing a frameshift mutation and a longer protein product (Yang et al. 2022). Grey European geese had an allele C fixed while white geese had a 1-bp deletion fixed (Yang et al. 2022). In humans, MLANA aka MART-1 (melanoma antigen recognized by T cells 1) forms a complex with PMEL17 (premelanosome protein, also known as SILV), affecting its expression, stability, trafficking, and processing (Hoashi et al. 2005). PMEL17 is needed for melanosome structuring and is a well-known colour dilution gene (Cieslak et al. 2011). In other avian species, mutation in *MLANA* causes a spotted Almond-phenotype in domestic pigeon (*Columba livia domestica*; Bruders et al. 2020) and *MLANA* is also differentially expressed in feather follicles of grey and white Chinese domestic geese (Zhao et al. 2025).

Buff is a rare colour (Fig. 2G, Table S1), occurring in certain goose breeds (Brecon Buff, Buff Back, American Buff and Suchovy goose). Buff is a sex-linked recessive trait, i.e. is inherited within the Z chromosome, with buff females inheriting one copy from the father, while buff males inherit a copy from both parents. The age of mutation is unknown, but the buff-coloured Brecon Buff breed was developed in 1929 by Rhys Llewellyn by purchasing buff-coloured geese from a flock of grey and white geese and crossing a buff female to a white Embden gander (Llewellyn 1934). Though more likely a non-spotting carrying gander of the type ‘almost white male’ as in the Pilgrim goose breed, was actually used due to the rapid establishment (4 years) of a flock breeding true to colour (Ashton 2021). Origin of the American Buff is undocumented (Ashton 2021), and it is unknown if the buff mutation in different breeds have independent origins. The genetic basis for buff is unknown, but in several birds, mutations in *TYRP1* (*tyrosinase related protein 1*) have caused brown, light brown or reddish plumage tone (Nadeau et al. 2007, Cortimiglia et al. 2016, Li et al. 2019, Ghosh Roy et al. 2025). TYRP1 is a key enzyme contributing to the melanin biosynthetic pathway, and *TYRP1* is a strong candidate gene for buff colour, as it is in the Z chromosome in geese, consistent with the inheritance patterns.

Blue is also a rarer colour in geese, mainly observed in Franconian, American and Faroese geese and the European and Chinese goose crossbreed Steinbacher (Fig. 2I, Table S1). Blue is an autosomal incompletely dominant trait (Ashton 2021; Table 1), and two copies of the blue allele result in silver/lavender colour (Fig. 2H-2I). The buff allele in combination with the blue allele creates lilac (heterozygous for blue) and cream (homozygous for blue) colours. The genetic basis for blue is unknown, and several genes could be causing blue.

**Table 1.**
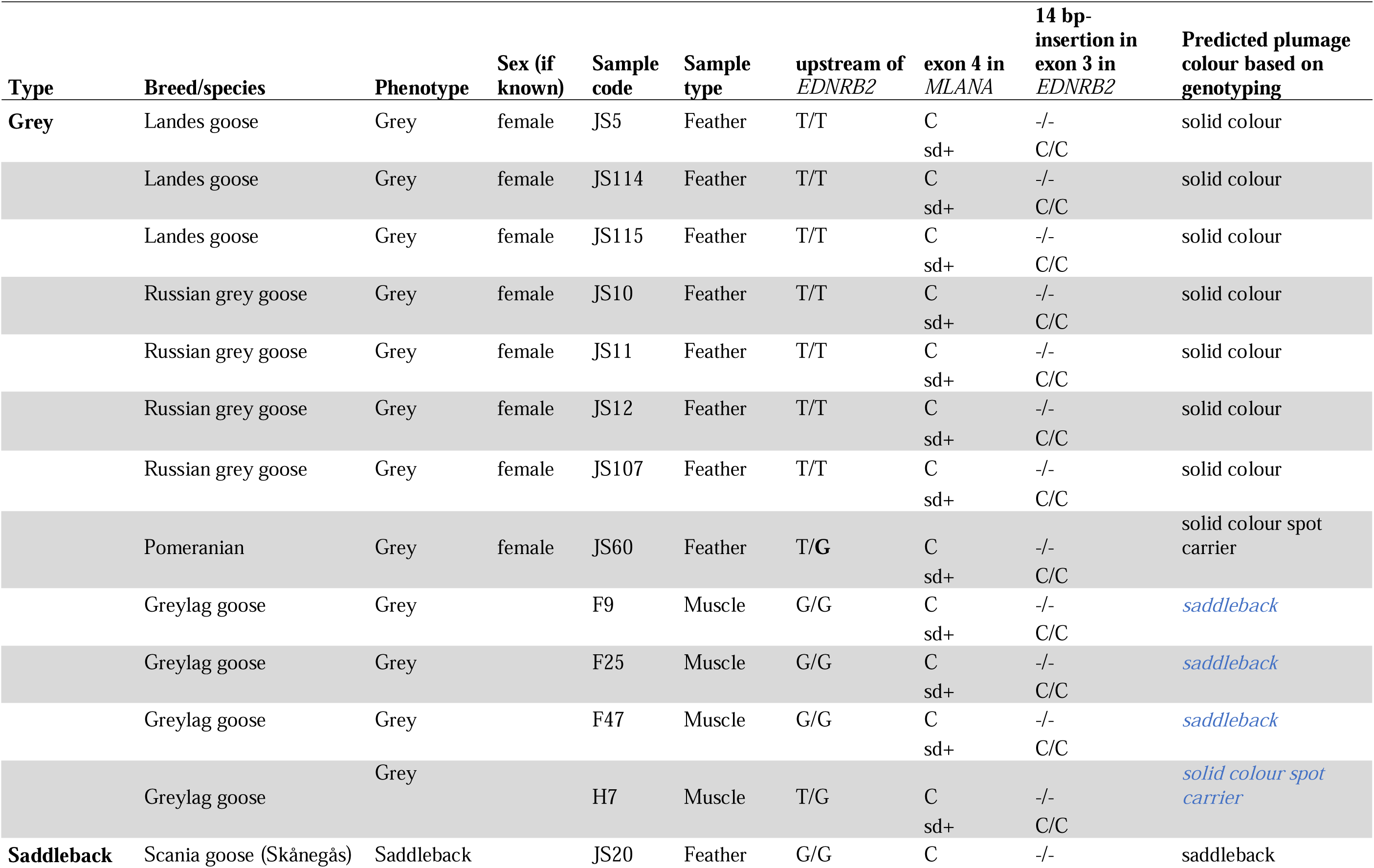
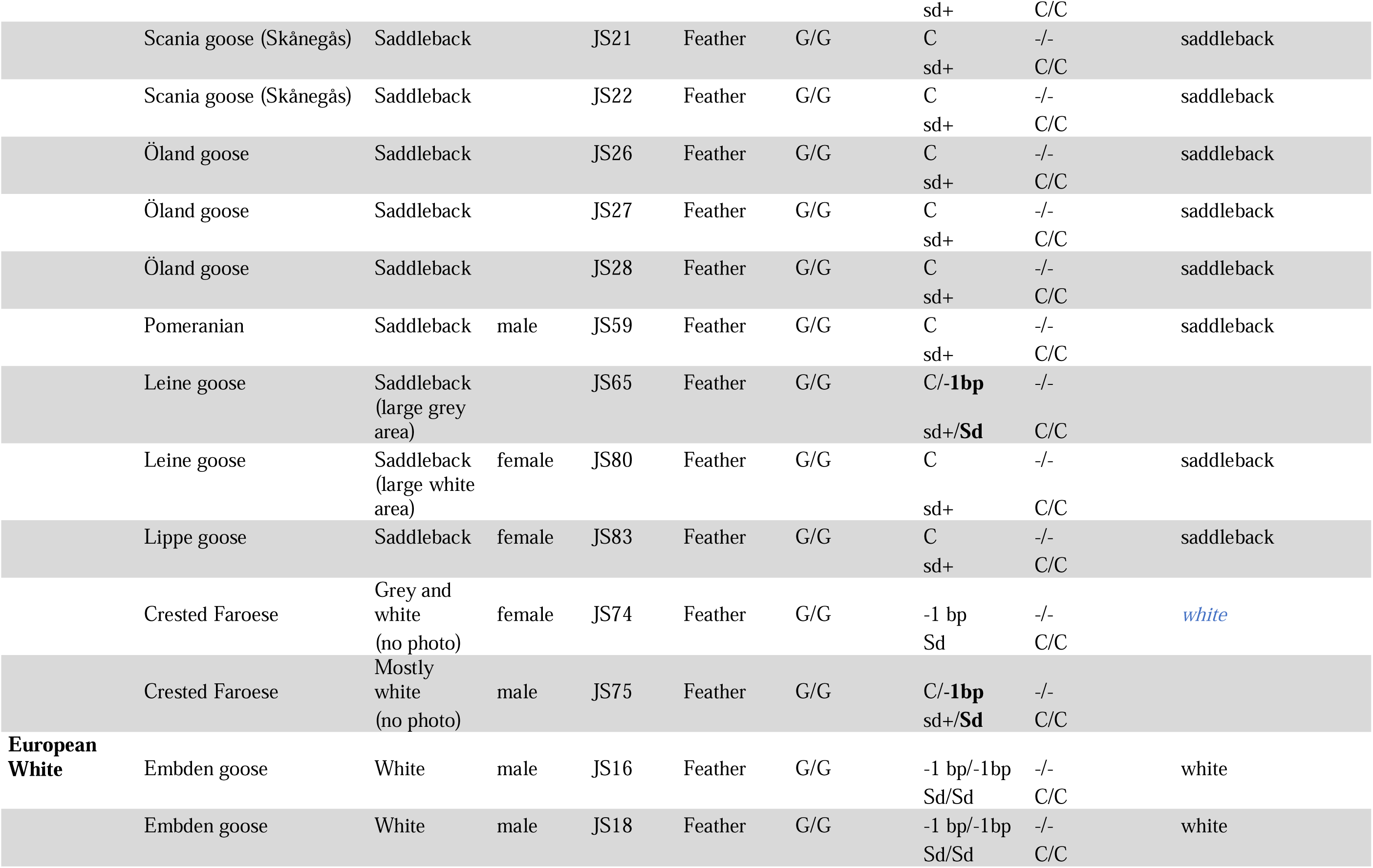
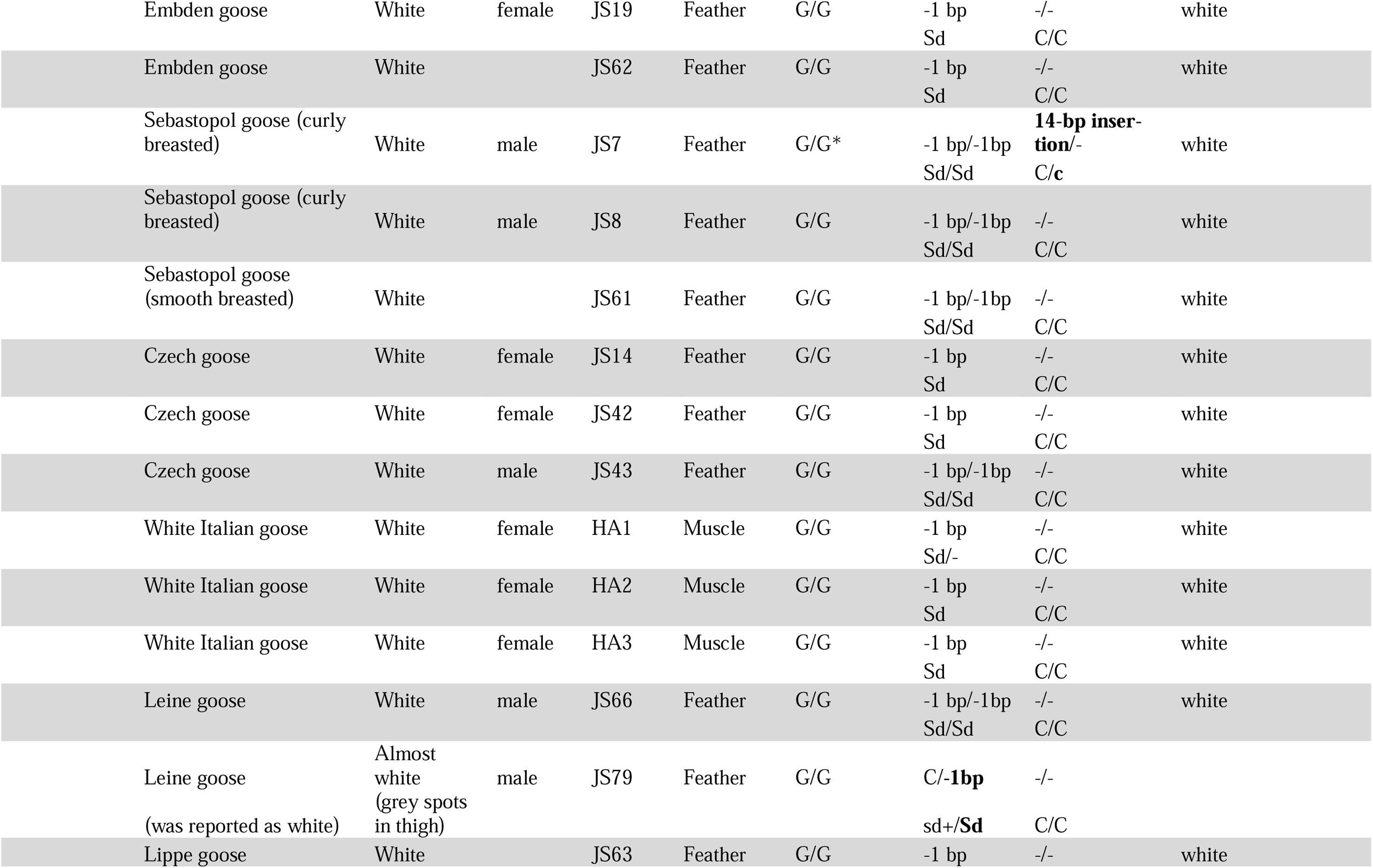
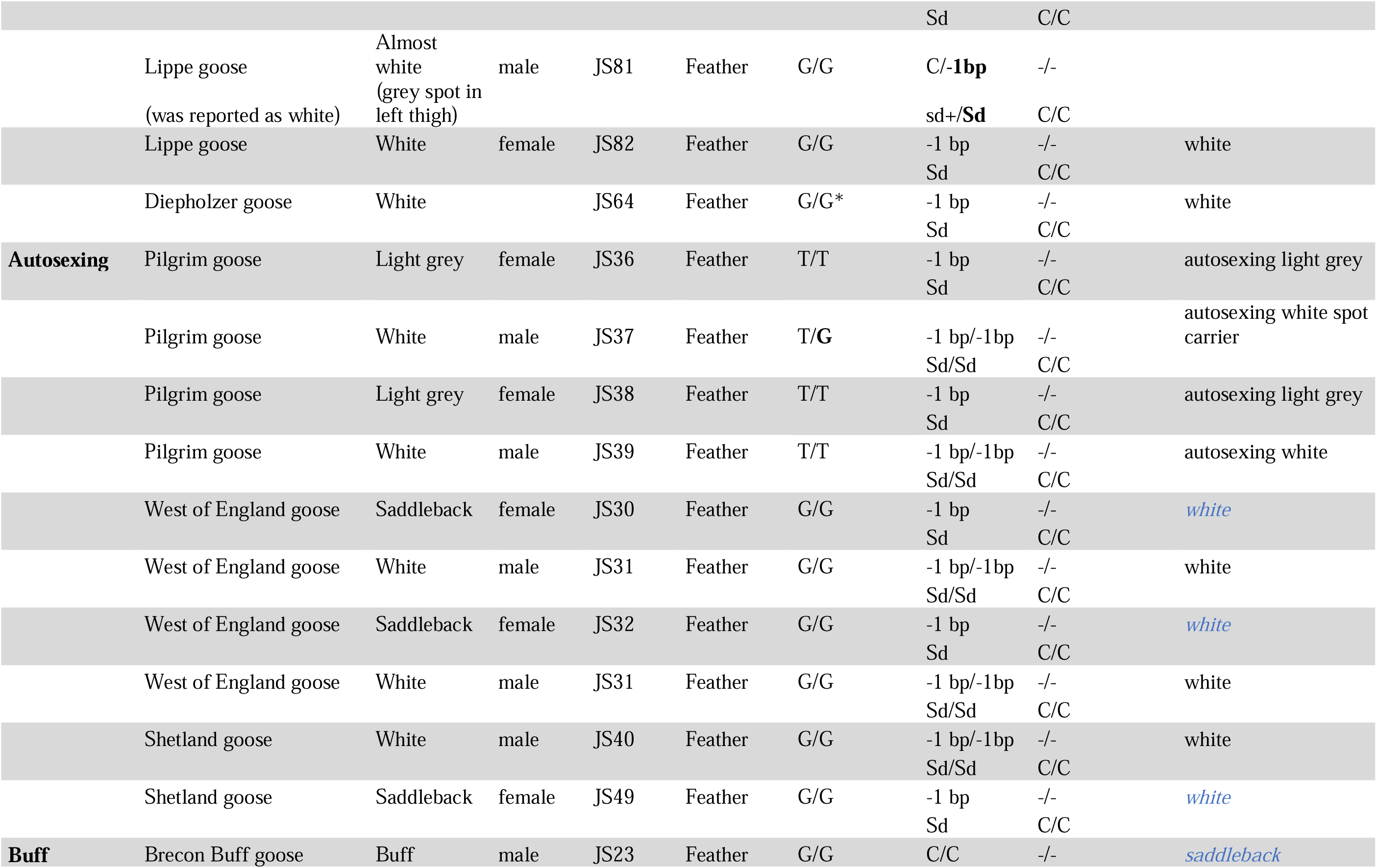
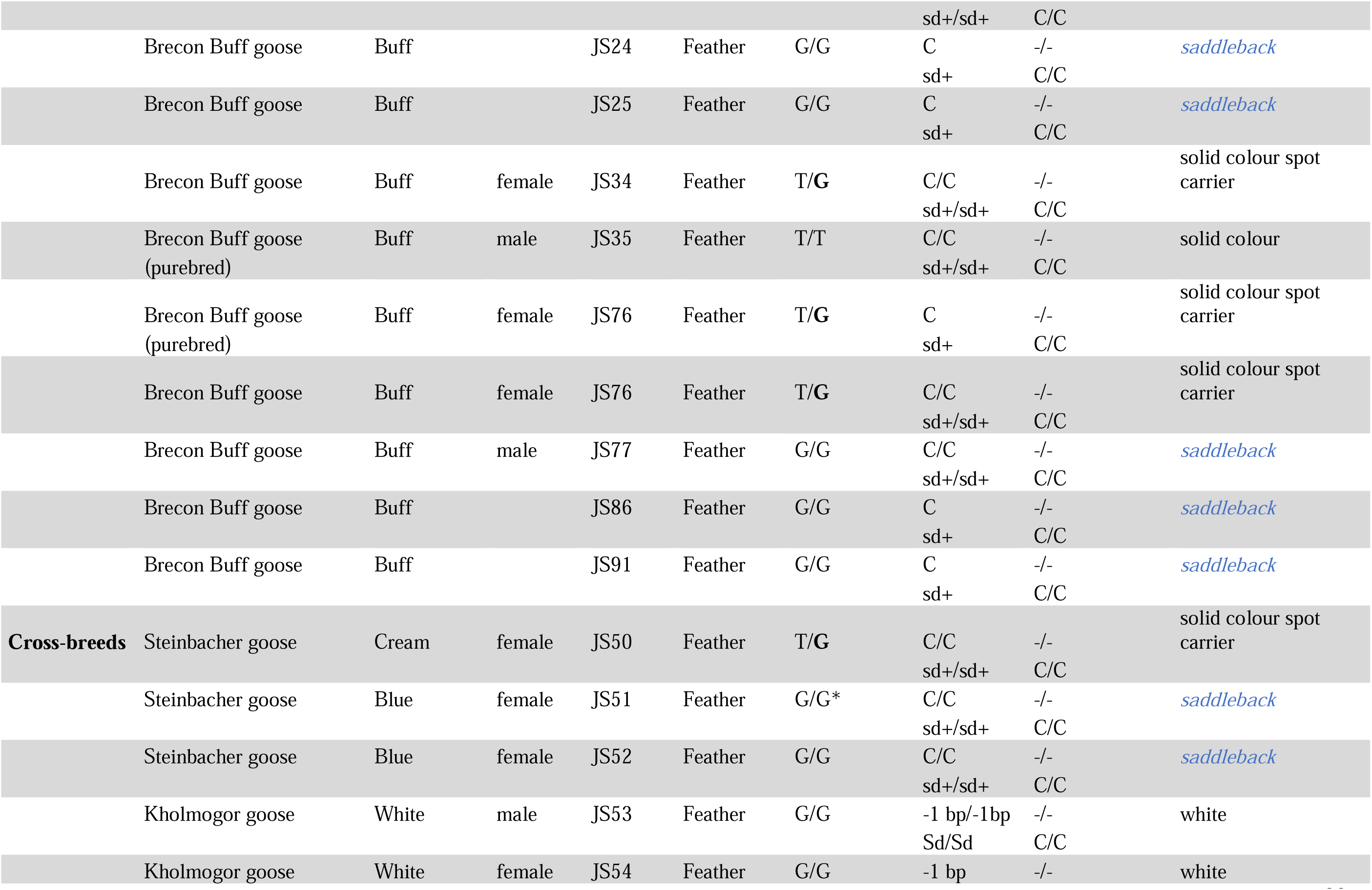
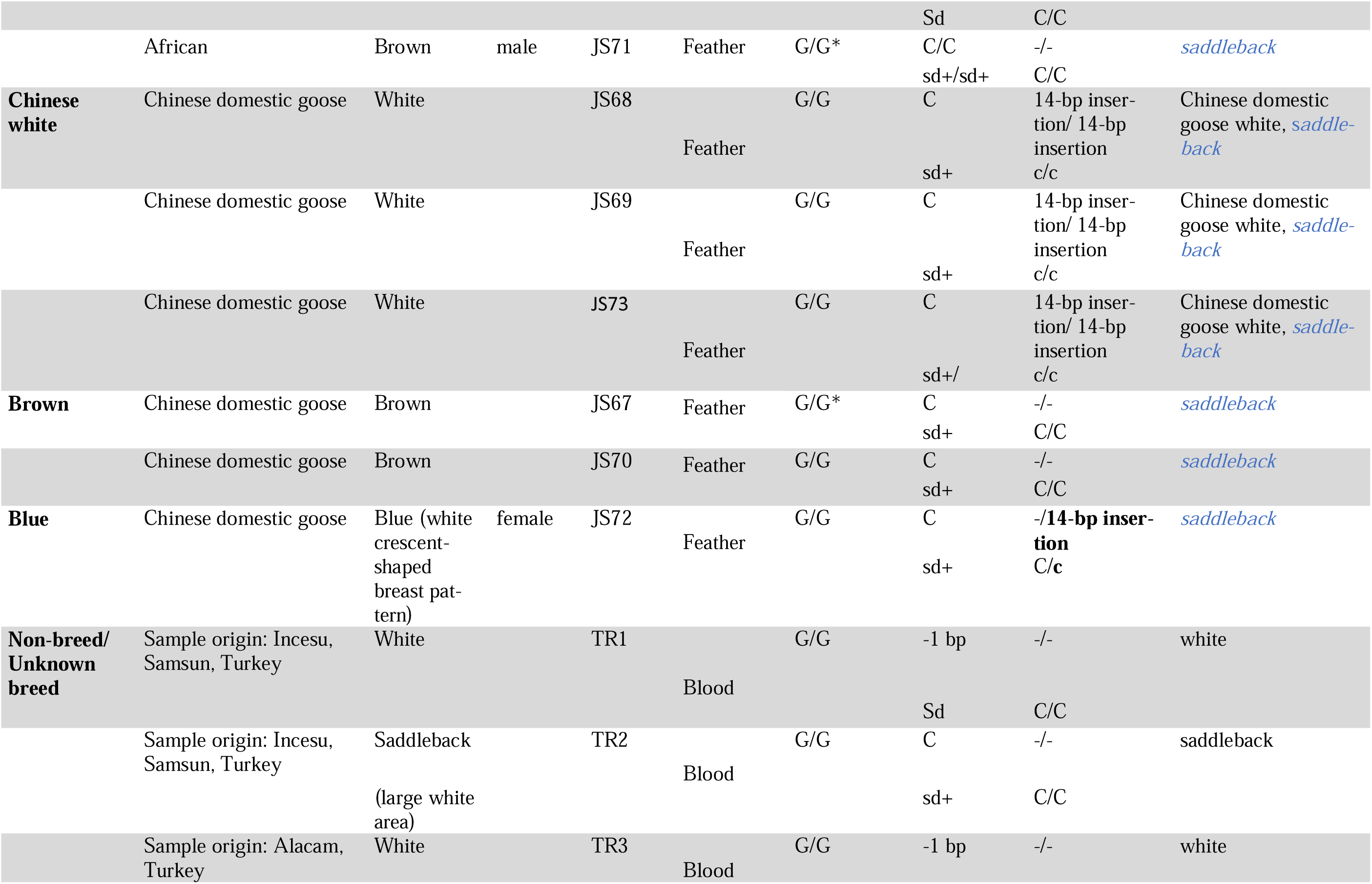
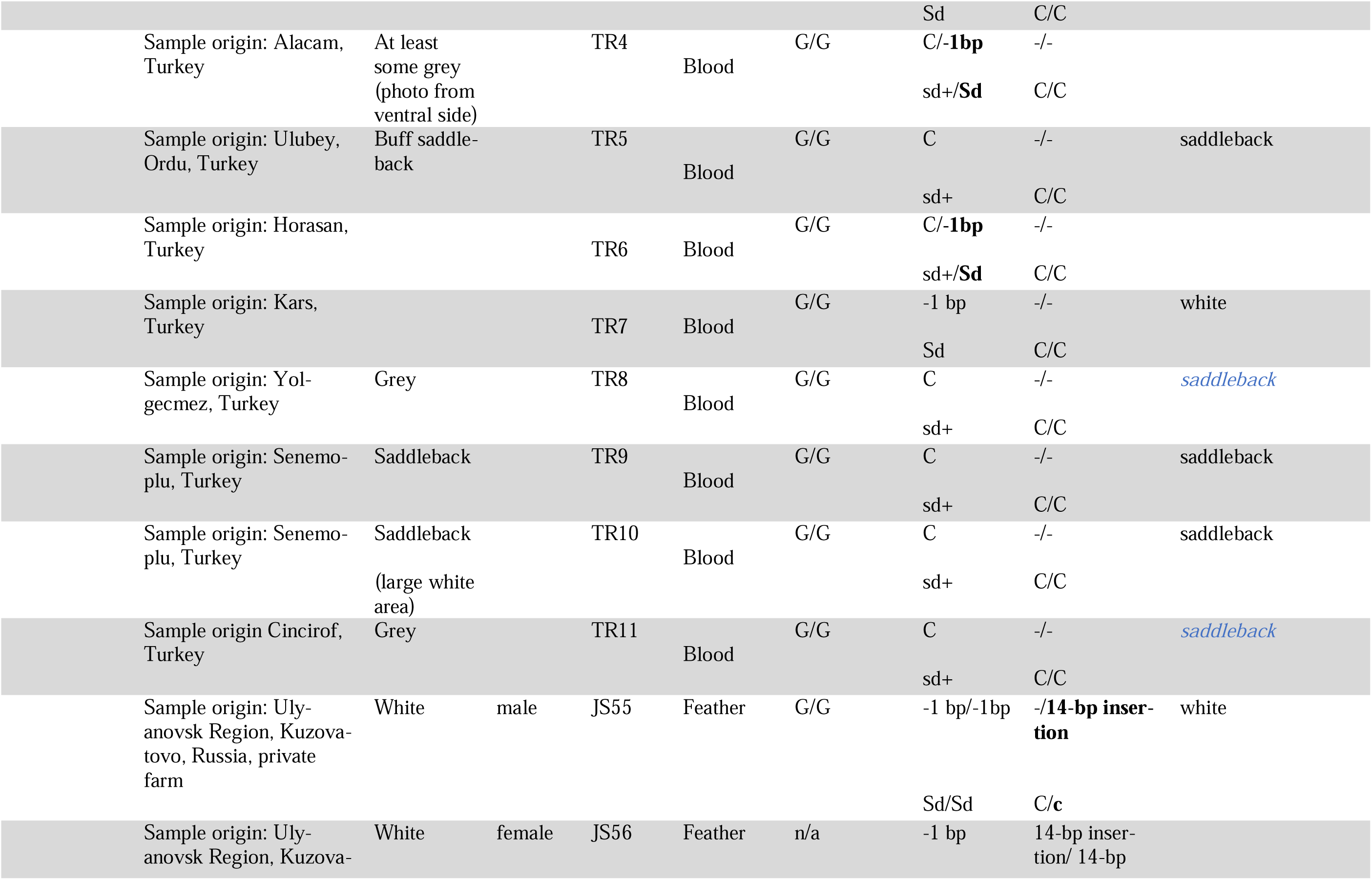
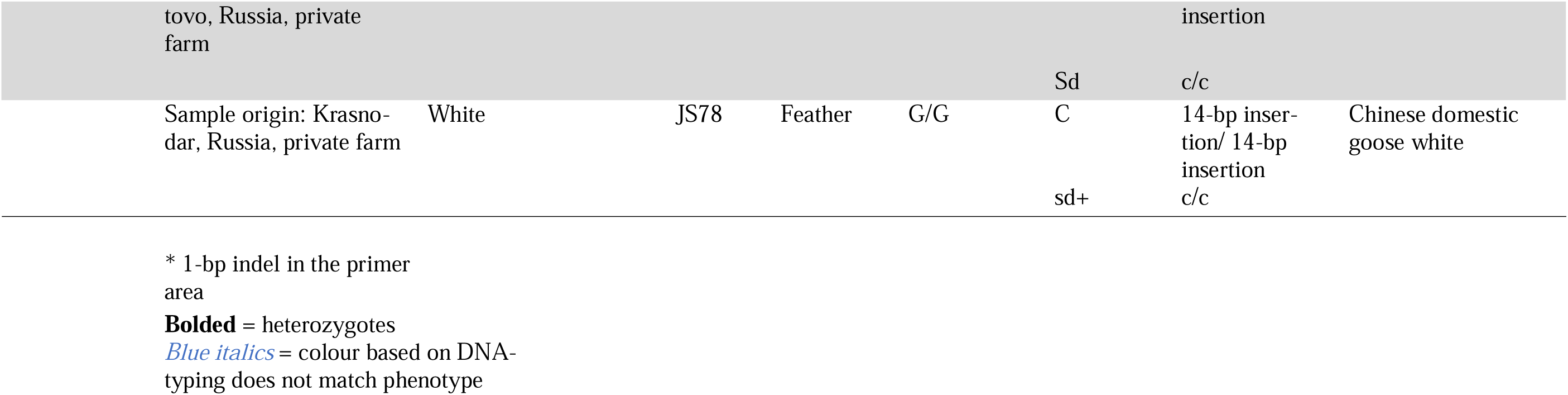
Genotypes of the studied domestic goose breeds (*Anser anser* -derived, *A. cygnoid* -derived and crossbreeds) as well as wild greylag geese. T = linked to solid colour, wild type, in domestic geese and G = linked to spotting in domestic geese in the locus upstream of *EDNRB2*. C = no dilution, wild type and - 1 bp = sex-linked dilution in *MLANA*. Due to sex-linkage in *MLANA*, sequencing cannot reveal if an individual is homozygous male or hemizygous female (except for heterozygous males, as both alleles manifest), unless the sex of the samples is known. In Chinese domestic geese - = no insertion, coloured/wild type and 14-bp insertion = white, non-colour. We also show the same genotyping results with common allele symbols for the mutation on *MLANA,* found to be in perfect association with the sex-linked dilution phenotype, and for Chinese domestic goose white: Sd = sex-linked dilution, sd+ = no sex-linked dilution, wild-type, C = coloured, wild type and c = white, non-colour (gosling yellow). n/a = sequencing failed. Bolding indicates a heterozygous individual, blue italics font the cases in which our expectations were not met and * indicates a 1-bp deletion in the primer area.

Two colour phenotypes are known for the Chinese domestic goose: white and wild type brown, also known as grey (Fig. 2K, 2M, Table S1). White colour in Chinese domestic goose is due to a 14-bp insertion (NW_013185915.1: g. 750,748–750,735 insertion) in exon 3 of the *EDNRB2* gene, though the white geese have small grey spots on the back (see Fig. 1b in Xi et al. 2020). This insertion causes a frameshift mutation in the *EDNRB2* gene, leading to a nonsense-mediated mRNA decay, causing a lack of melanin pigment (Xi et al. 2020). Geese homozygous for the 14-bp insertion are white (Xi et al. 2020, Yang et al. 2022, Ouyang et al. 2022, Yang et al. 2024), while geese heterozygous for the 14-bp insertion display a white breast pattern of variable size (Fig. 2L), which can extend to the dorsal side and white primary flight feathers (Yang et al. 2024). This indicates codominance of the white and brown plumage colours, although this mutation can have different penetrance or other loci involved (Yang et al. 2024). However, Chinese domestic geese can also have the blue phenotype (Fig. 2J, 2L), and the blue allele is also found in the crossbreed African goose. It is not certain if the mutation was introgressed from European geese to Chinese geese or vice versa.

The GWAS identified candidate mutations behind the white plumage colour in European domestic goose need to be verified using a larger sample size and geese of different colours, as only two breeds were used in the original study. We aimed to genotype the two mutations; the upstream mutation of the *EDNRB2* gene (NW_013185915.1: g. 775,151 G > T) and the 1-bp deletion in exon 4 of the sex-linked *MLANA* gene (NW_013185876.1: g.950,868 C > -) in different goose colour phenotypes (grey, white, saddleback, buff, blue and both types of autosexing) and in the greylag goose, to verify if these mutations are causative. As the GWAS identified candidate mutation for spotting was tens of thousands of bases upstream of the transcription start site of the *EDNRB2* gene, we also searched for the causative mutation in the coding region of the *EDNRB2* and its downstream *VAMP7* gene. Additionally, we genotyped the 14-bp insertion causing white colour in Chinese domestic geese to check for crossbreeding between European and Chinese domestic geese, as these forms can hybridize. Our assumptions of the expected genotypes in differently coloured European and Chinese domestic geese are presented schematically in Fig. 4. Using candidate gene approach, we further sequenced the coding sequence of the *TYRP1* gene, as a candidate for the buff colour.

**Fig. 4.**
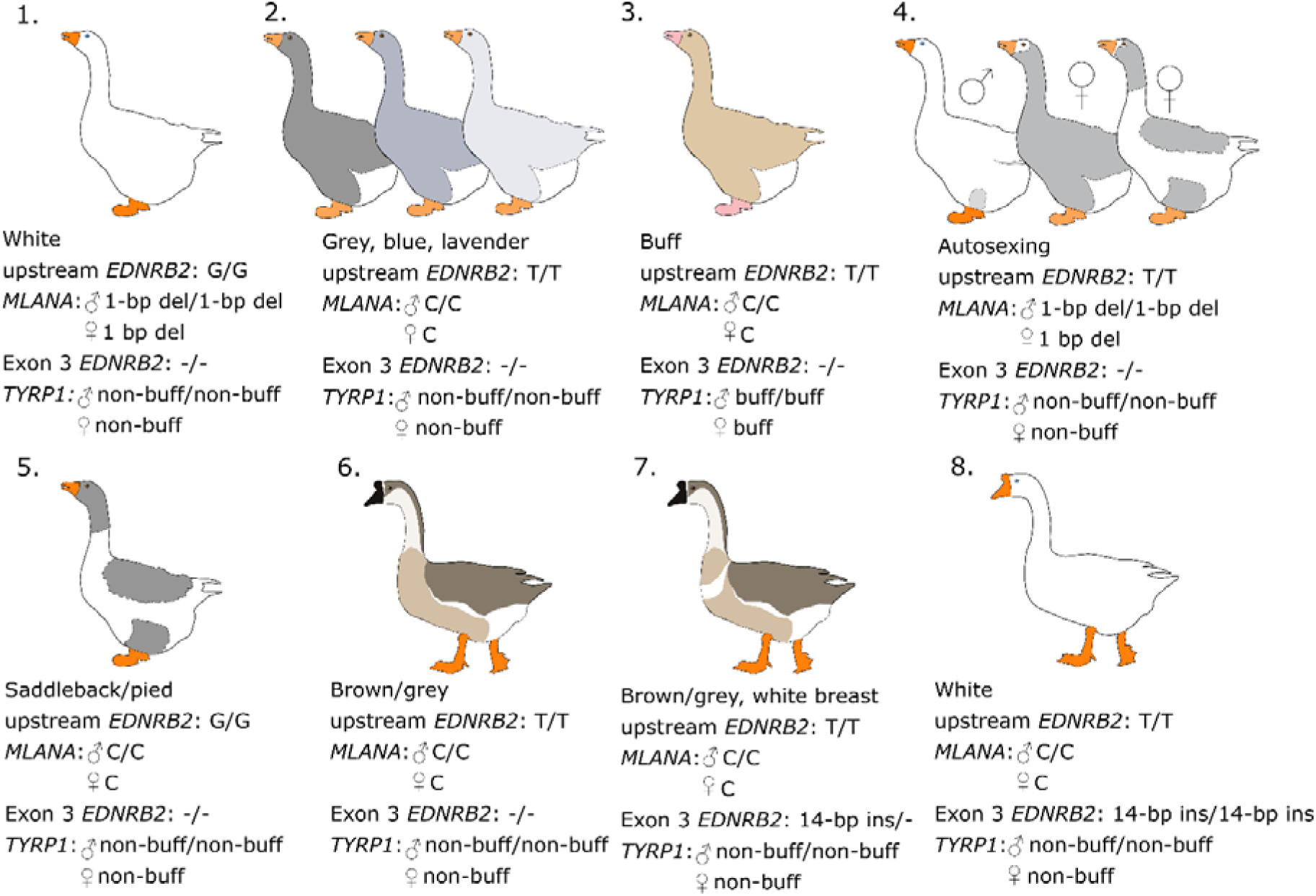
Schematic drawings of predicted genotypes of European domestic geese (*Anser anser* -derived) [1-5] and Chinese domestic geese (*A. cygnoid* -derived) [6-8] based on previous knowledge. 1. White goose homozygous for nucleotide G (assumed spotting allele) upstream of the *EDNRB2* gene and homozygous/hemizygous for the 1-bp deletion within the *MLANA* gene (assumed sex-linked dilution allele). 2. Grey and blue breeds and wild greylag geese are expected to be wild type in both loci (T/T in upstream of *EDNRB2* and no deletions in *MLANA*). 3. Buff geese are expected to also have the same genotype as grey geese in these two loci, but we expect to find a mutation in the coding sequence of the *TYRP1* gene. 4. Autosexing breeds are expected to be homozygous/hemizygous for the 1-bp deletion within the *MLANA* (sex-linked dilution). 5. Saddleback breeds are expected to be homozygous for nucleotide G (spotting). 6. Wild type, i.e. brown Chinese domestic geese, are expected to have a similar genotype as greylag geese in these loci. 7. Chinese domestic geese heterozygous for the 14-bp insertion are expected to have a white breast pattern, and 8. white Chinese domestic geese are homozygous for the 14-bp insertion within the exon 3 of the *EDNRB2* gene, while European breeds should not carry this insertion. The size of breeds in the schematic drawings is not in real scale as the sizes of different breeds vary.

## Materials and methods

### Sampling and DNA extraction

Our samples consisted of different breeds of the European domestic goose (*n* = 59) and non-breed geese from Turkey and Russia (*n* = 14), greylag geese (*n* = 4), crossbreeds of European and Chinese domestic geese (*n* = 6) and Chinese domestic geese (*n* = 6), altogether 89 samples (Table 1). Additionally, eight greylag geese and three bean geese (*A. fabalis*) were used in some analyses (*n* = 11, see below). Most colour phenotypes (grey, European white, saddleback, two types of autosexing, buff, blue, cream and Chinese white, brown and blue) were included except lilac and silver. Samples were donated by private individuals, a goose farm, zoos keeping domestic geese (Moscow Zoo, Zoo Neuwied, Tierpark Hamm, and Arche Warder, with support from the Association of Zoological Gardens [Verband der Zoologischen Gärten (VdZ) e.V]) or hunters. Forty-four individuals were previously analysed in Heikkinen et al. (2015, 2020), of which 27 were re-extracted in this study. DNA from feathers (*n* = 73) was extracted as in Honka et al. (2022), DNA from muscle tissue (*n* = 4) was extracted using E.Z.N.A. Tissue DNA Kit (Omega Bio-Tek) according to the manufacturer’s protocol and DNA from bone samples (*n* = 6) was extracted as in Appendix S2 of Kvist et al. (2022).

### Genotyping *EDNRB2* and *MLANA* mutations

We designed primers flanking the candidate mutations upstream of *EDNRB2* and *MLANA* genes and flanking the 14-bp insertion within the *EDNRB2* gene (Table S2) based on Chinese domestic goose whole-genome sequence (NW_013185915.1). PCR reactions for *MLANA* and *EDNRB2* 14-bp insertion were performed in 10 _μ_l volumes using 1 x Phusion HF buffer (Thermo Fisher Scientific), 0.2 mM of each dNTPs, 0.5 _μ_M of F- and R-primers (Table S2), 0.02 U/_μ_l of Phusion DNA Polymerase (Thermo Fisher Scientific), and 1 _μ_l of template DNA, which was diluted to 25 ng/µl for muscle tissue samples. The thermocycling conditions were 98°C for 4 min, followed by 40 cycles of 98°C for 30 s, 60°C for 30 s for *MLANA* and 63 °C for 30s for *EDNRB2*, and 72°C for 40 s with a final extension of 72°C for 7 min. The PCR reactions for the mutation upstream of *EDNRB2* were performed in 10 µl volumes using 1 x QIAGEN Multiplex PCR Master Mix, 0.2 µM of F- and R-primers (Table S2) and RNase-free water. The thermocycling conditions were 95 °C for 15 min, followed by 40 cycles of 94 °C for 30 s, 63 °C for 90 s and 72 °C for 90 s with a final extension of 72 °C for 10 min. Seven of the greylag goose samples and three bean goose samples were analysed only for the upstream *EDNRB2* mutation (Table S3). All samples were sequenced in one direction using the F-primers with BigDye Terminator v.3.1 (Applied Biosystems). The sequencing reactions were run on an ABI3730 Genetic Analyzer (Applied Biosystems), and the alleles were checked visually using the program CodonCode Aligner v.4.0.4. (CodonCode Corporation) and recorded.

### Long-range sequencing of *EDNRB2* and *VAMP7* genes

To search for the causative locus of spotting in coding regions of the adjacent genes *EDNRB2* and *VAMP7*, we PCR amplified the genes and used Oxford Nanopore’s MiniON sequencing device to sequence the PCR products (except the last exon of *VAMP7,* which we Sanger-sequenced). We designed four overlapping primer pairs (Table S2) to amplify the whole *EDNRB2* gene and around 25,000-bp region upstream of the gene (altogether ∼37,000 bp). Primers were designed based on whole-genome sequences from two Chinese domestic goose (NW_025927715.1 and NW_013185915.1) and other closely related species: duck (*Anas platyrhynchos;* NC_051781.1), tufted duck (*Aythya fuligula*; NC_045571.1), ruddy duck (*Oxyura jamaicensis*; NC_048896.1), black swan (*Cygnus atratus*; NC_066374.1), mute swan (*C. olor*; NC_049181.1), and chicken (*Gallus gallus*; NC_052535.1). We designed primers for the upstream and downstream regulatory regions or introns of the *VAMP7* gene, but no DNA of the correct size was amplified (results not shown), and hence the primers lie within exons (5’ and 3’ untranslated regions) flanking the coding sequence (CDS) (Table S2). The last 27 bp are, however, missing from exon 6 in *VAMP7*. Primer design was based on the whole-genome sequence of Chinese domestic goose (NC_089885.1) and other closely related species: duck (NC_051781.1) and mute swan (NC_049181.1).

We performed long-range PCR for *EDNRB2* and its upstream regulatory region in four separate PCR reactions (∼10,000 bp in each reaction) and in one reaction for the *VAMP7* (8,866 bp) for the first six exons for muscle tissue samples. The PCR reactions were performed in 12.5 µl volumes using 1 x Q5® High-Fidelity Master Mix (NEB), 0.5 µM of F- and R-primers (Table S2) and RNAse-free water. The thermocycling conditions were 98 °C for 30 s, followed by 35 cycles of 98 °C for 30 s, primer-dependent annealing temperature (Table S2) for 30 s and 72 °C for 8 min 20 s with a final extension of 72 °C for 2 min. The amplification was successful for four white domestic goose and two greylag goose muscle samples for *EDNRB2* and for three white domestic goose and one greylag goose muscle sample for *VAMP7*. To sequence the amplified products, sequencing libraries were prepared for white domestic and greylag goose separately with the Native Barcoding Kit 96 V14 (SQK-NBD114.96) (Oxford Nanopore Technologies, ONT) according to the manufacturer’s protocol, except for incubating the samples for 10 min at 65°C and then at fridge (+4 °C) overnight during native barcode ligation before pooling samples. This modification was done to avoid barcode jumping, and the prepared libraries were sequenced on the MinION MK1C on a R10.4.1 flow cell. The raw data (fast5) was basecalled and demultiplexed using Dorado Basecaller, version [1.2.0] (Oxford Nanopore Technologies), in super accuracy mode. The reads were aligned to a reference Chinese domestic goose (NW_013185915.1) using minimap2 (Li, 2018, 2021) and samtools (Danecek et al., 2021). Even though overnight incubation in the fridge was performed, some barcode jumping was observed, and we treated the domestic goose library as a pool of white geese and the greylag goose library as a pool of wild type individuals. The alignments of domestic and wild individuals were visually checked to find candidate positions for the causative mutation.

The last (7^th^) exon of *VAMP7* (161 bp) was amplified with a similar PCR reaction and reaction conditions (primers in Table S2) as the upstream of *EDNRB2* mutation (see above), except the annealing temperature was 60 °C. The amplification was successful for four white domestic goose and three greylag goose muscle samples. It was also similarly Sanger-sequenced as the upstream *EDNRB2* mutation and *MLANA* SNP in both directions using the PCR primers.

### Sequencing coding regions of *TYRP1*

We designed primers to amplify the exons of *TYRP1* gene based on sequences of Chinese domestic geese (NC_089912.1 and KR080366). Each exon was amplified in a separate PCR reaction, except the first longer exon (372 bp) was amplified with two overlapping primer sets, as feather samples can have low-quality DNA with shorter fragment lengths. We used four buff, four grey, four white and four saddleback geese feather samples. PCR reactions were performed similarly to the mutation upstream of the *EDNRB2* gene and within the *MLANA* gene (see above), except for different primers (primers in Table S1). Reaction conditions were similar as well, except the annealing temperatures differed between the primer pairs, see Table S2. Similarly, the PCR products were Sanger-sequenced in both directions.

## Results

### Genotyping *EDNRB2* and *MLANA* mutations

We observed that the upstream of the *EDNRB2* mutation (NW_013185915.1 g. 775,151 G > T) was not causative of the spotting, as all solid-colour greylag geese, bean geese and Chinese domestic geese were homozygous for the G allele, except three greylag geese (Table 1). Two greylag geese were genotype T/T and one heterozygous T/G, hence both alleles are segregating in greylag goose populations (Table 1, Table S3). However, all individuals of grey domestic goose breeds (Landes and Russian grey) had T/T genotypes (Table 1). One grey Pomeranian (JS60) was heterozygous T/G and likely a solid colour spot carrier, which is not surprising as Pomeranian geese have both saddleback and grey (solid) colours. None of the grey geese (domestic or wild) carried the 1-bp deletion in the *MLANA* gene (NW_013185876.1: g.950,868 C > -), as predicted.

All individuals of saddleback-coloured breeds (Scania and Öland goose) and saddleback Pomeranian and Lippe goose showed expected genotypes: G/G in the upstream of the *EDNRB2* and wild-type in *MLANA* (Table 1), though Lippe goose is regarded as a white breed. Individuals of white breeds (Embden, Sebastopol, Czech, White Italian and Diepholzer) showed the anticipated genotypes G/G in the upstream of the *EDNRB2* and 1-bp deletion in *MLANA* (homozygous or hemizygous) (Table 1). Czech geese, which have been reported to be of “lighter” gosling plumage colour than other white breeds, carried the same sex-linked dilution alleles as other white breeds, however, other unknown loci could be involved. One Lippe individual had the anticipated genotype for a white breed (JS82), but another reportedly white male Lippe (JS81) was heterozygous for the sex-linked dilution, i.e. carried the 1-bp deletion in *MLANA* in a heterozygous state, while being homozygous G/G in the upstream of the *EDNRB2*. Upon re-examination, the gander was found to have a grey spot on the thigh.

A German heritage breed, the Leine goose, can have white, saddleback or grey colour. One saddleback Leine goose individual (JS65) with an extensive grey saddleback had genotype G/G upstream of the *EDNRB2* while being heterozygous for the 1-bp deletion in *MLANA.* This indicates a male individual (only males can be heterozygous), but was reported as female. The male partner of this goose (JS66) was white with the expected white genotype (G/G in the upstream of the *EDNRB2* and homozygous for the 1-bp deletion in *MLANA*). In another Leine pair, the female (JS80) was saddleback with a large white area (Fig. 5), carrying the expected genotype (G/G in the upstream of the *EDNRB2* and wild type in *MLANA*). Its male partner (JS79) was reported as white but found to be heterozygous for the 1-bp deletion, while being homozygous G/G in the upstream of the *EDNRB2*. Also in this case, a closer inspection of the gander revealed a light grey speckling on the thigh (Fig. 5). Similarly, two non-breed Turkish domestic geese (TR4 and TR6) were heterozygous for the 1-bp deletion in *MLANA*, while having G/G genotype in the upstream of the *EDNRB2.* The first individual has a grey head, but due to a photo being taken from the ventral side, other colouration is not visible. The second was a gosling with saddleback pattern in the fluff.

**Fig. 5.**
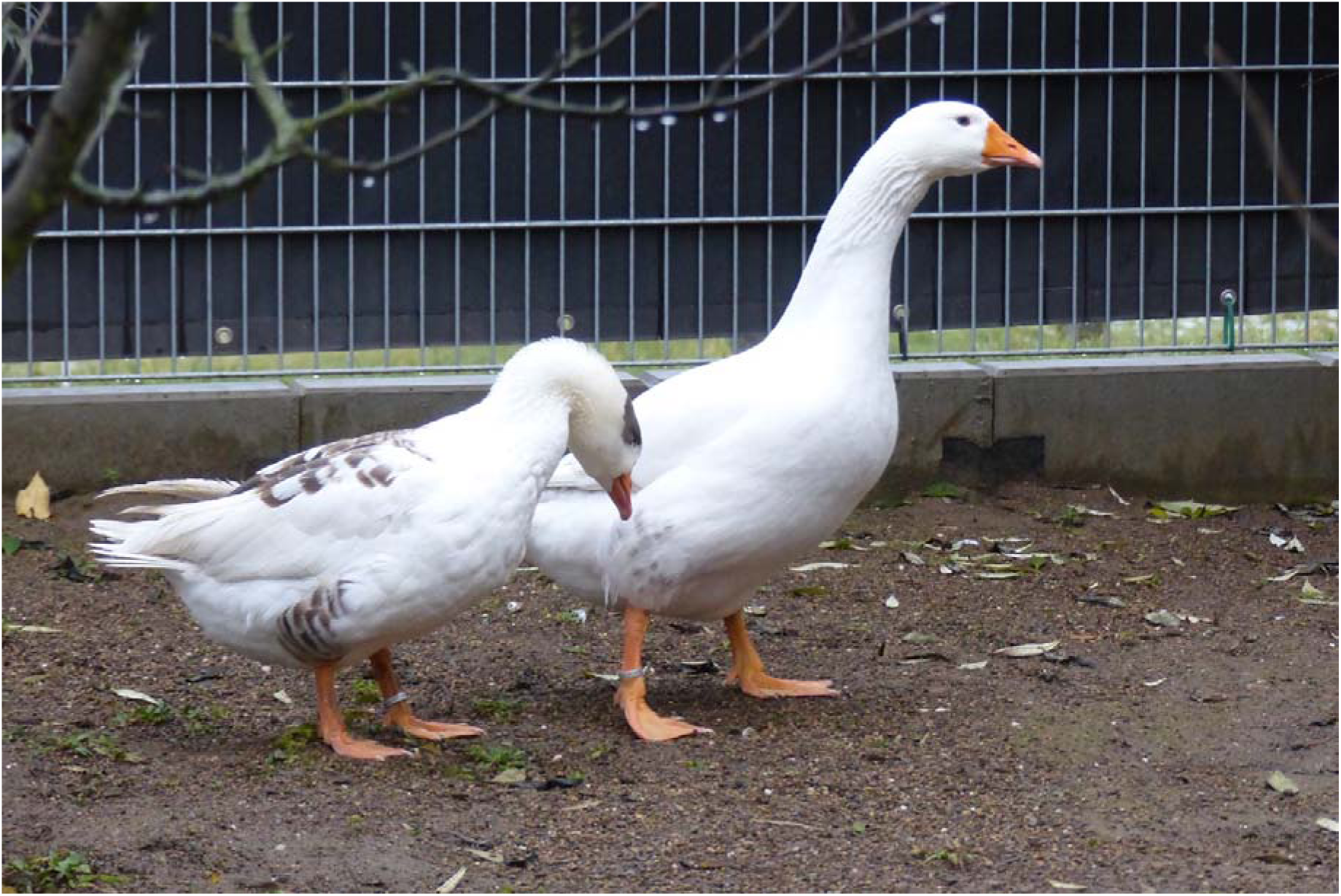
Leine goose pair (*Anser anser*-ancestry), with the female on the left and the male on the right. The female is homozygous for a mutation linked to spotting upstream of the *EDNRB2* gene, showing large white areas in plumage but does not carry sex-linked dilution (wild type). The male is also homozygous for the mutation linked to spotting upstream of *EDNRB2*, but heterozygous for the sex-linked dilution mutation in the *MLANA* gene. This individual was reported as white, but later inspection revealed that it has light grey speckling in the thigh coverts. Photo: Maximilian Birkendorf, Zoo Neuwied

As predicted, autosexing geese with almost white male and light grey female (e.g. Pilgrim breed) were homozygous for wild type allele in the presumed solid colour locus (T/T in the upstream of the *EDNRB2*) and homozygous/hemizygous for the 1-bp deletion in *MLANA*, except one Pilgrim male (JS37) was heterozygous (T/G) and thus presumed spot carrier (Table 1). This individual had one white flight feather in his first and second years, which indicates it being phenotypically also a spot carrier (non-spot carriers have some grey in flight feathers). West of England and Shetland geese present the second type of autosexing, in which the male is almost white and the female saddleback. Individuals of these breeds were homozygous G/G upstream of *EDNRB2* and homozygous/hemizygous for the 1-bp deletion in *MLANA.* Based on our prediction, this genotype should produce white males and females. This also confirms that another unknown sex-linked locus is involved in this form of autosexing (Ashton 2021).

Two Crested Faroese geese were genotyped, with a saddleback Crested Faroese female (JS74) found to be G/G in the upstream of the *EDNRB2* and having the 1-bp deletion in *MLANA*, thus genotypically like the white breeds. It is possible that the West of England/Shetland/Normandy/Bavent goose type autosexing (almost white male and saddleback female) is also present in the Faroese goose population, due to their variable plumage patterns. White Crested Faroese male (JS75) was genotype G/G upstream of the *EDNRB2* while being heterozygous for the 1-bp deletion in *MLANA.* As this sample was collected 14 years ago, we did not have access to a photo to verify if the plumage was completely white.

Out of ten Brecon Buff geese, six were homozygous G/G in the locus upstream of the *EDNRB2* gene, and two were heterozygous (T/G), while none carried the 1-bp deletion in *MLANA* (Table 1). Based on our predictions, the six individuals with G/G genotype should be spotted, but Brecon Buffs are a solid-coloured breed. Crossbreeds between European domestic and Chinese domestic goose (Steinbacher and African goose), as well as brown and blue geese also had unexpectedly G/G in the locus upstream of the *EDNRB2* gene, but no 1-bp deletion in *MLANA* (Table 1). One cream coloured Steinbacher (JS50) was heterozygous T/G in the locus upstream of the *EDNRB2.* Also, grey non-breed Turkish domestic geese were homozygous G/G in the locus upstream of the *EDNRB2.* These results also confirm that the G/G genotype is not causative of the spotting.

All white Chinese domestic geese carried the 14-bp insertion in the exon 3 of the *EDNRB2* gene in a homozygous state as expected (Table 1). Additionally, one blue Chinese domestic goose individual (JS72) carried one copy of the 14-bp insertion (heterozygous) and phenotypically had the white crescent shape breast pattern that is typical for heterozygotes (Table S4). Two geese of unknown breed from Russia were homozygous for the 14-bp insertion, while one was heterozygous. Brown Chinese domestic geese did not carry the 14-bp insertion as predicted. One European domestic goose individual of the curly-breasted Sebastopol breed (JS7) carried the 14-bp insertion in a heterozygous state.

The results of the genotyping of the mutation upstream of *EDNRB2* gene, the 1-bp deletion in *MLANA* gene and the 14-bp insertion in *EDNRB2* gene are summarised in Fig. 6. Additionally, phenotypes of each genotype are presented schematically.

**Fig. 6.**
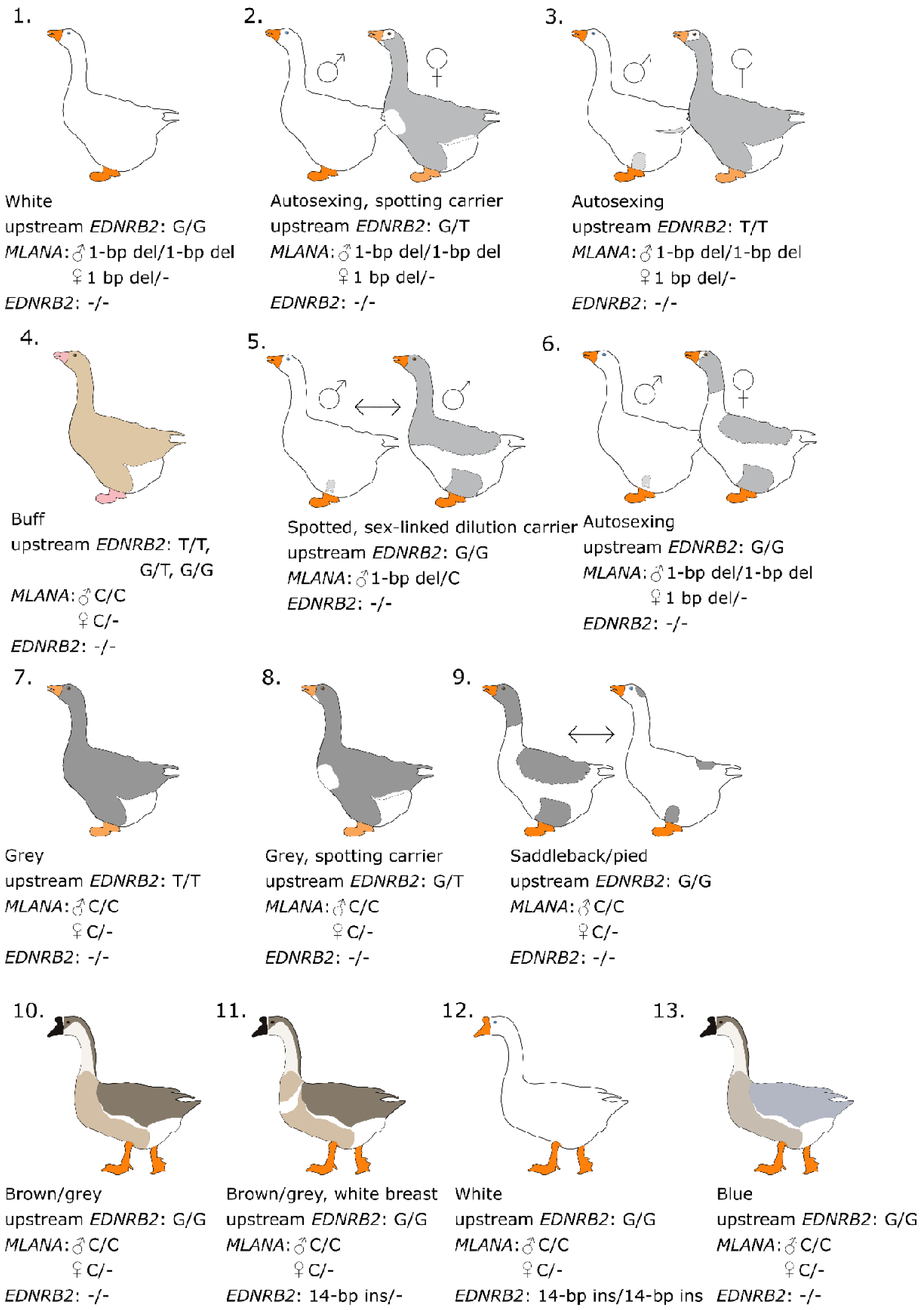
Genotypes of European domestic geese (*Anser anser* -derived) [1-9] and Chinese domestic geese (*A. cygnoid* -derived) [10-13] in three loci genotyped in this study and a simplified schematic drawing of the phenotype associated with the genotype. 1. White European goose, 2. autosexing spot-carrying (heterozygous for spotting) goose with an almost white male (whiter than non-spot carrying individual) and light grey female with a white chin, a white breast batch and/or white flight feathers, 3. autosexing goose with an almost white male and light grey female, often with white forehead and white feathers around the eyes known as ‘spectacles’, 4. buff goose, 5. saddleback male heterozygous for sex-linked dilution (dilution carriers showed variable expressivity from small grey spotting on rump to larger grey areas), 6. autosexing goose with an almost white male and saddleback female, another locus seemingly allows underlying saddleback to be visible in females despite being homozygous for sex-linked dilution, 7. grey goose, 8. grey goose heterozygous for spotting (spot carriers show a white chin, a white breast batch and/or white flight feathers), 9. saddleback, the pattern has variable expressivity or another gene is affecting the amount of white areas, the pattern colour can be grey, buff, blue or lavender/silver, 10. brown/grey Chinese domestic goose, 11. brown/grey Chinese domestic goose with white breast pattern and/or white primary flight feathers (heterozygous for Chinese domestic goose white), the pattern shows variable expressivity as the size of the breast patch varies and it can extend to the dorsal size, 12. white Chinese domestic goose (homozygous for Chinese domestic goose white), 13. blue Chinese domestic goose. The size of breeds in the schematic drawings is not in real scale as the sizes of different breeds vary.

### Long-range sequencing of *EDNRB2* and *VAMP7* genes

We visually inspected the sequences of the domestic and greylag geese against the reference Chinese domestic goose sequences. We searched for sites in which the white domestic goose would differ from both the greylag and Chinese domestic goose, as both are non-spotted, i.e. solid colour. We found potential variation associated with colour in two intronic regions and in one indel upstream of the *EDNRB2* gene. We designed primers to amplify these variations in additional goose individuals, and Sanger sequenced these loci, but no association with colour was found (results not shown). No such variation was found in the coding sequence of the *VAMP7* gene either. Hence, we did not proceed to genotype more samples.

### Sequencing coding regions of *TYRP1*

After visually inspecting the sequences of the domestic and greylag geese, we found no sequence variation in the coding sequence of the *TYRP1* gene that was associated with feather colour. No further genotyping was thus performed.

## Discussion

We confirmed the mutation identified by Yang et al. (2022) in the *MLANA* gene (NW_013185876.1: g.950,868 C > -) to be causative of sex-linked dilution in the European domestic goose, and contributing to white plumage colouration. White plumage in the European domestic goose is caused by two independent genes (Jerome 1953), but we could not identify the causative mutation for the spotting, which, together with the 1-bp deletion in the *MLANA* gene, causes white. Mutation 18,846 kb upstream of the predicted transcription site of the *EDNRB2/LOC106047519* gene, identified as a strong candidate mutation by Yang et al. (2022), was found not to be the causative, as most greylag geese, buff-coloured Brecon Buff geese, and all Chinese domestic geese and bean geese had the assumed “spotting” genotype, even though they are solid coloured.

G/G genotype upstream of *EDNRB2* (NW_013185915.1: g. 775,151 G > T) seems to be, however, strongly linked to spotting in domestic geese except for Brecon Buff geese and non-breed geese as individuals in grey domestic goose breeds were homozygous for the T/T genotype, while all individuals of white breeds and saddleback breeds were homozygous for the G/G genotype. Due to the strong linkage, we searched for the causative mutation from the *EDNRB2* gene and regulatory regions upstream of it, as well as from the coding sequence of the *VAMP7* gene. This gene is the adjacent gene downstream of the *EDNRB2,* and due to low genetic diversity of European domestic geese (Heikkinen et al. 2020), the linked region could be longer. No variation associated with colour was found in either of the genes, though we were unable to sequence the regulatory region of the *VAMP7* gene. We attempted to design primers extending beyond the exons, but apparently, the regions were too variable compared to the Chinese domestic goose. An annotated genome for the greylag goose unfortunately does not currently exist. Based on our study, the *EDNRB2* gene or the coding region of the *VAMP7* gene are not causative of the spotting, and further genomic work is needed to elucidate the genetic background of spotting, such as GWAS analyses. Similarly, further work is required to elucidate if other loci are involved in controlling the extent of white spotting, which can vary from saddleback pattern (Fig. 2C) to minimal grey markings on the head and back (Fig. 5, Table S4). Small-scale breeding experiments have also pointed to the possibility of another locus being involved in controlling the amount of white patterning (Ashton & Ashton 2022, Ashton & Ashton 2023).

Similarly, we sequenced the exons in the *TYRP1* gene, a strong candidate gene for buff colour, but found no variation associated with colour. We only focused on sequencing exons, thus, it is possible that the causative mutation lies in the regulatory region or intron splicing sites. We conclude that the coding region of the *TYRP1* is not causative of buff colour and further genomic work is warranted.

Additionally, we found that the two types of autosexing, ‘almost white male and light grey female’ and ‘almost white male and saddleback female’, might have different genetic backgrounds. Both types exhibited the sex-linked dilution (homozygous in males with two Z chromosomes and hemizygous in females with one Z chromosome) as predicted, but ‘almost white male and light grey female’ type expressed T/T genotype upstream of the *EDNRB2* gene linked to wild type phenotype. ‘Almost white male and saddleback female’ type expressed the G/G genotype linked to spotting upstream of the *EDNRB2* gene. However, white geese similarly have G/G upstream of the *EDNRB2* gene and 1-bp deletion in the *MLANA* gene. Thus, autosexing geese with the ‘almost white male and saddleback female’ colouration probably have another likely sex-linked locus affecting the colour (Ashton 2021) and further work is also warranted to elucidate the genetic background.

We also noted one Pilgrim goose individual carrying the G allele linked to spotting upstream of the *EDNRB2* and thus carrying spotting, which was also then confirmed phenotypically. In this autosexing breed the spotting is unwanted and spot-carrying heterozygous Pilgrim males can be identified due to their whiter appearance and pink blood quills on the wing tips at the age of 4-5 weeks, and heterozygous females by a white chin spot, a white breast patch and/or white flight feathers (C. & M. Ashton, personal communication, Table S4). Purebred Pilgrim males have purple flight feather quills around the age of 5 weeks, diluted grey primary and secondary flight feathers as adults, and diluted grey on rump (C. & M. Ashton, personal communication, Table S4). Even though the spotting locus was not identified in this study, the linked mutation upstream of the *EDNRB2* could possibly be used to help eliminate the spotting in breeds in which it is unwanted.

To our knowledge, we are the first to describe sex-linked dilution heterozygotes in association with probable spotting, though the number of such individuals was low. Two such individuals were reported as phenotypically white, but on closer inspection, they were found to have grey or light grey speckling in thigh coverts (Fig. 5.). One individual was saddleback with an extensive grey area in the back, though the sex of this individual remained unclear, and one had at least grey in the head. Based on these observations, the expressivity of the sex-linked dilution may vary or other loci might be involved. More heterozygous individuals should be studied to determine the range of expressivity. This information could be useful to goose breeders aiming for pure flocks.

We found one individual in the Sebastopol breed to be heterozygous for the 14-bp insertion in the *EDNRB* gene causative of white plumage colour in the Chinese domestic goose (Table 1), thus indicating Chinese domestic goose introgression in this breed. The UK Sebastopols were crossed with Chinese domestic geese in the 1970s (C. & M. Ashton, personal communication), and this ancestry is also visible in genome-wide analyses performed on the same individuals of this study (Heikkinen et al. 2020). The crossbreeding background was additionally observed in Chen et al. (2023) with an independent sample set.

To conclude, we confirmed that the 1-bp deletion in exon 4 of the *MLANA* gene identified by Yang et al. (2022) is causative of the sex-linked dilution colour and responsible for autosexing in geese, in which the male is almost white and the female light grey. Causative mutation for spotting was not identified, but the G to T mutation upstream of the *EDNRB2* gene was found to be in strong linkage to solid (grey) colour in European domestic geese, except in the Brecon Buff breed. Each genotype and predicted phenotype based on genotyping of the *MLANA* and *EDNRB2* mutations are presented in Table S4, based on current best knowledge of goose colour inheritance. Further work is required to elucidate the causative mutation of spotting and thus the white colour, which is caused by both the sex-linked dilution and spotting. We also showed that the Sebastopol breed has been crossbred with the Chinese domestic goose, due to the presence of the 14-bp insertion in exon 3 of the *EDNRB2* gene, known as the causative mutation for white colour in Chinese domestic geese.

## Acknowledgments

We thank Virpi Rantalainen and Hauhalan Hanhifarmi for muscle tissue samples, Chris and Mike Ashton, N.A.R Wood, Denise Moss, M. Reeves, C. J. Boothby, John Burns, S. Pavlova and the Moscow Zoo, Mikael Olsson, Kirsi Mäki, Brigitte Rohrhuber and Tierpark Ueckermünde, Stella Pontius and Zoo Hellabrunn, Maximilian Birkendorf and Zoo Neuwied, Felicitas Schulte and Tierpark Hamm, Sinje Büttner, Anabell Jandowsky and Arche Warder, for feather samples. In addition, we thank Julia Kögler and the Association of Zoological Gardens and Gerrit Wehrenberg for help in obtaining samples. We are grateful to Vesa-Matti Murtomäki for providing a greylag goose tissue sample. We thank Tuula Pudas and the Zoological Museum of the University of Oulu for providing bone samples. We also thank Driss Eddahbi for help with DNA extractions. We especially thank Chris and Mike Ashton for sharing knowledge on goose colour inheritance and phenotypes.

## Data availability

DNA sequences are either too short to be uploaded to GenBank (< 200 bp), are non-continuous (e,g, exons of *TYRP1* with intron sequences missing) or samples were treated as a sequence pool for long-range sequencing, thus, individual sequences cannot be distinguished. Sequences are available from the corresponding author upon reasonable request. Descriptions of predicted phenotypes of each genotypic combination genotyped in this study are described in Table S4.

## Funding

This research was funded by Biodiverse Anthropocenes Research Programme of the University of Oulu PROFI6 funding supported by the Research Council of Finland to JH.

## Conflict of interests

The authors declare that they have no conflict of interest.

## Ethics approval

Samples were either extracted in a previous study, obtained from legally hunted geese or from museums. Feather samples were collected by the bird owners, with appropriate import permits for samples outside of the European Union.

## Notes

### Competing Interest Statement

The authors have declared no competing interest.

