## Supplemental Table 4 for "Sex-linked dilution colour in the European domestic goose confirmed to be a 1-bp deletion in the *Melan-A* gene"

**Table S4**. Genotypes in two loci: upstream of *endothelin receptor B-like* (*EDNRB2*) and in exon 3 of the *melan-A* (*MLANA*) gene. One-base pair deletion in *MLANA* was confirmed in this study to be causative of sex-linked dilution, due to perfect association in domestic geese of greylag goose origin (*Anser anser*). Mutation upstream of *EDNRB2* was found to be linked to spotting in European domestic geese (except in buff-coloured geese), but not the causative mutation.

| **Upstream of *EDNRB2*, linked to spotting in European domestic geese, position: NW_013185915.1: g. 775,151 G > T** | ***MLANA,* sex-linked dilution, position: NW_013185876.1: g.950,868 C > -** | **Phenotype** | **Photo** |
| --- | --- | --- | --- |
| T/T | ♂ C/C or ♀ C  sd+/sd+ or -/sd+ | grey, solid colour | 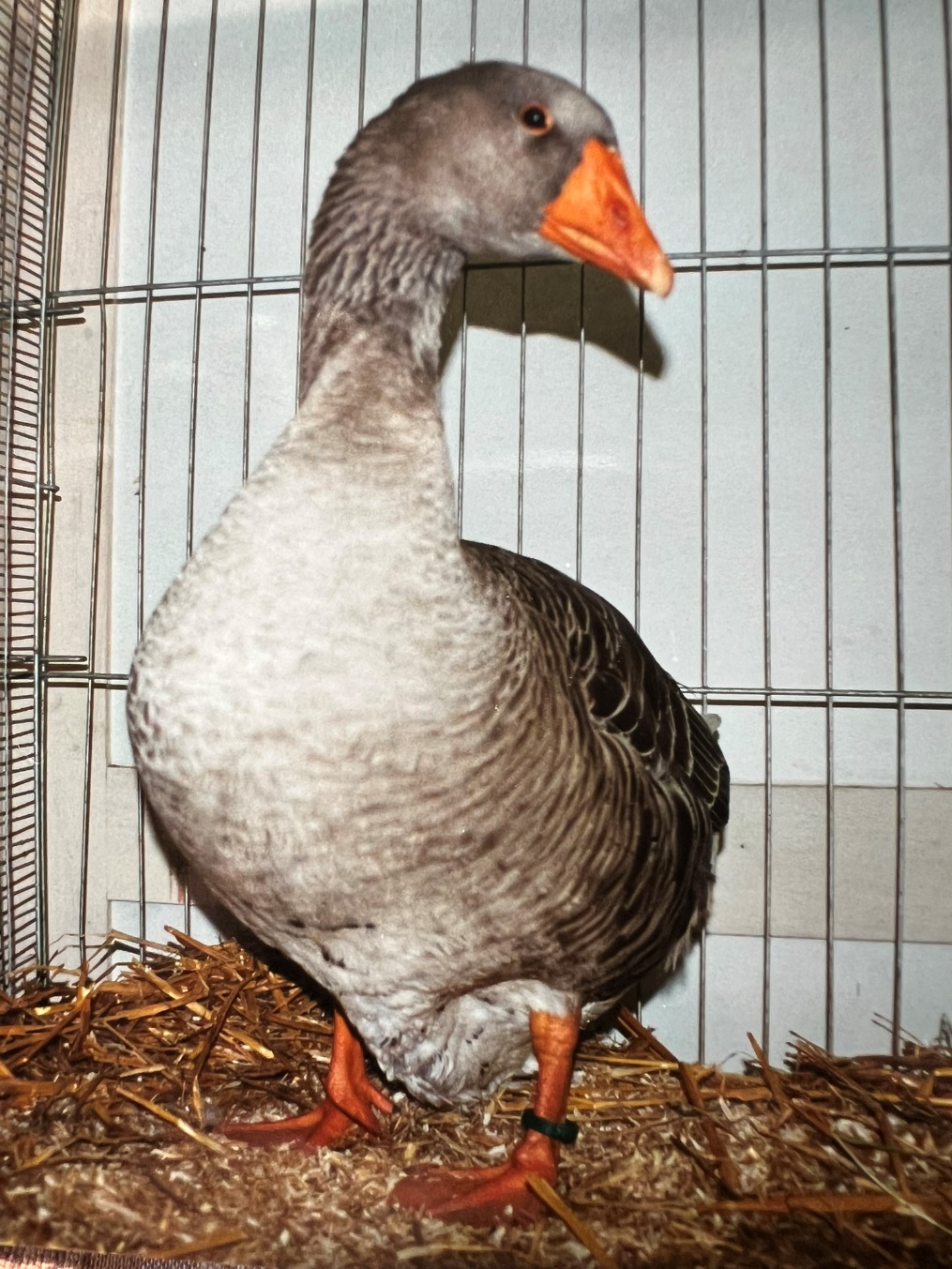  Pomeranian goose. Photo: Chris & Mike Ashton |
| T/G | ♂C/C or ♀ C  sd+/sd+ or -/sd+ | grey with white chin spot/white breast patch/white flights (T/G is very variable in its expression) | Photo not available |
| T/T | ♂ -1 bp/-1 bp  Sd/Sd | autosexing almost white male, e.g. Pilgrim breed (some grey in rump, wings and tail, purple primary quills at 4-5 weeks of age) | 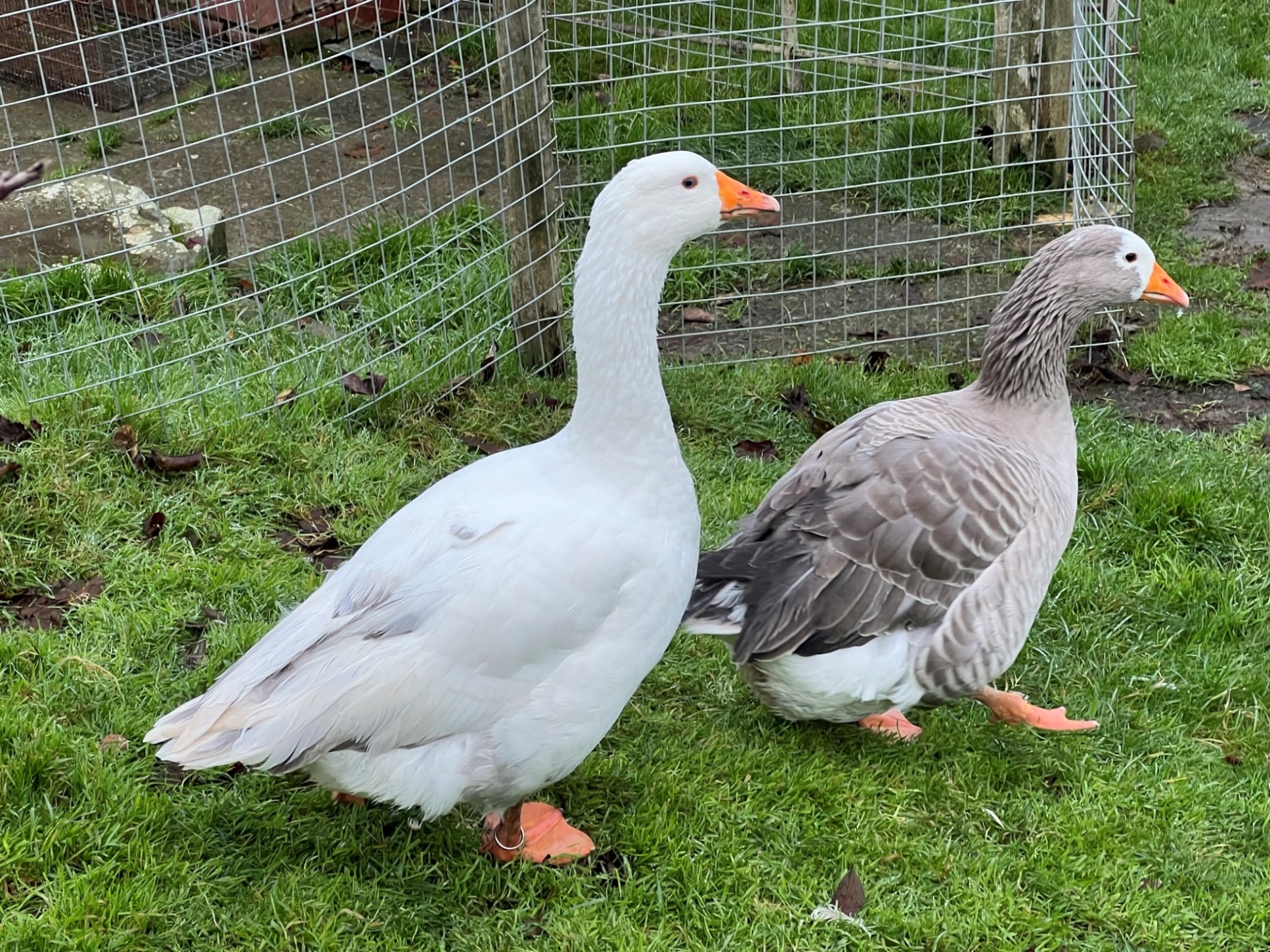  Pilgrim male. Photo: Chris & Mike Ashton |
| T/T | ♀ -1 bp  -/Sd | autosexing light grey female, often white around the eyes and forehead (so called spectacles). As the geese get older, they lose grey pigment from the face and can develop white feathers around the neck. E.g. Pilgrim breed | 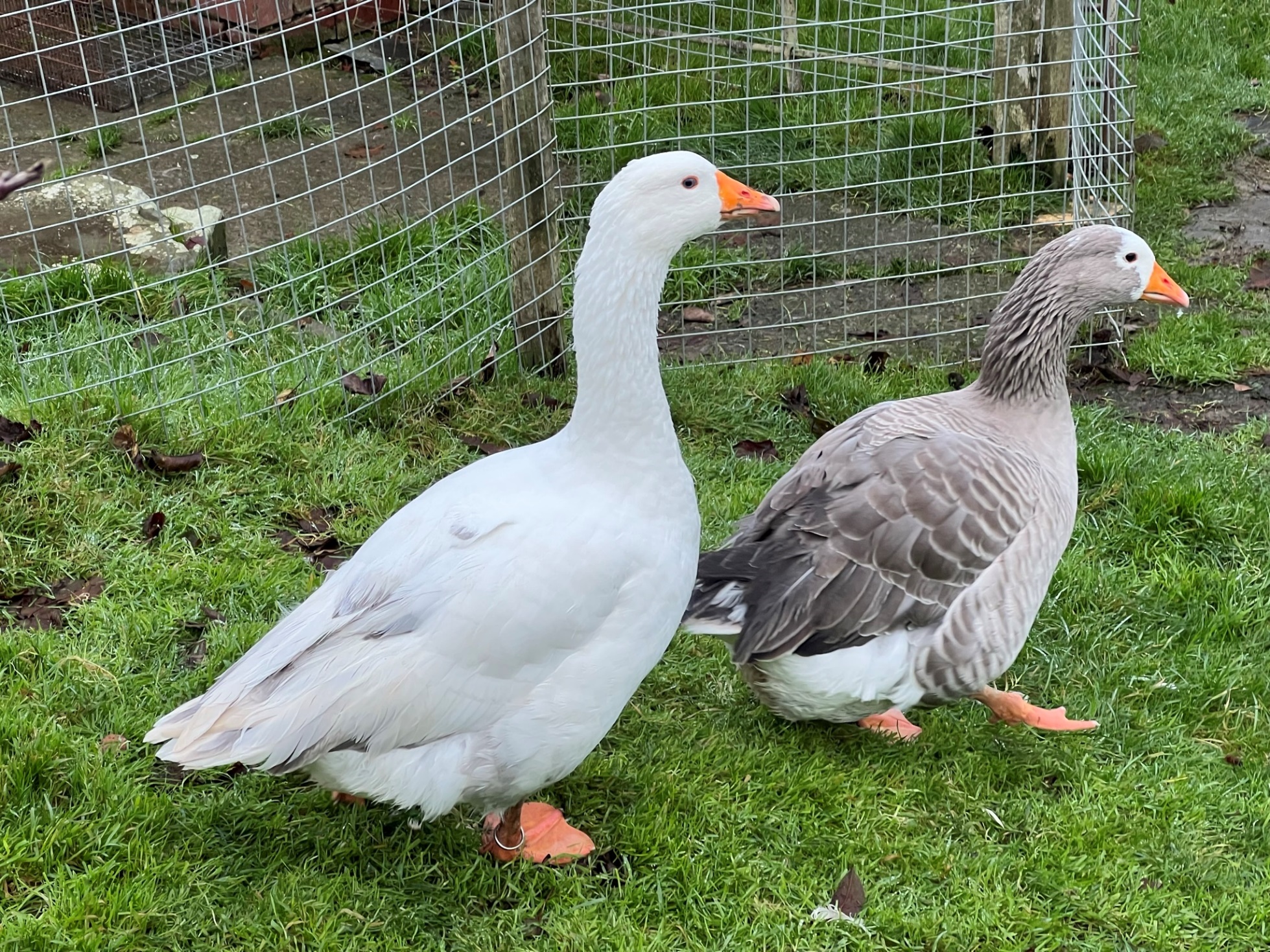  Pilgrim female. Photo: Chris & Mike Ashton |
| T/T | ♂ -1 bp/C  Sd/sd+ | light grey male but darker than light grey autosexing female (e.g. in Pilgrim breed, see above photo) | 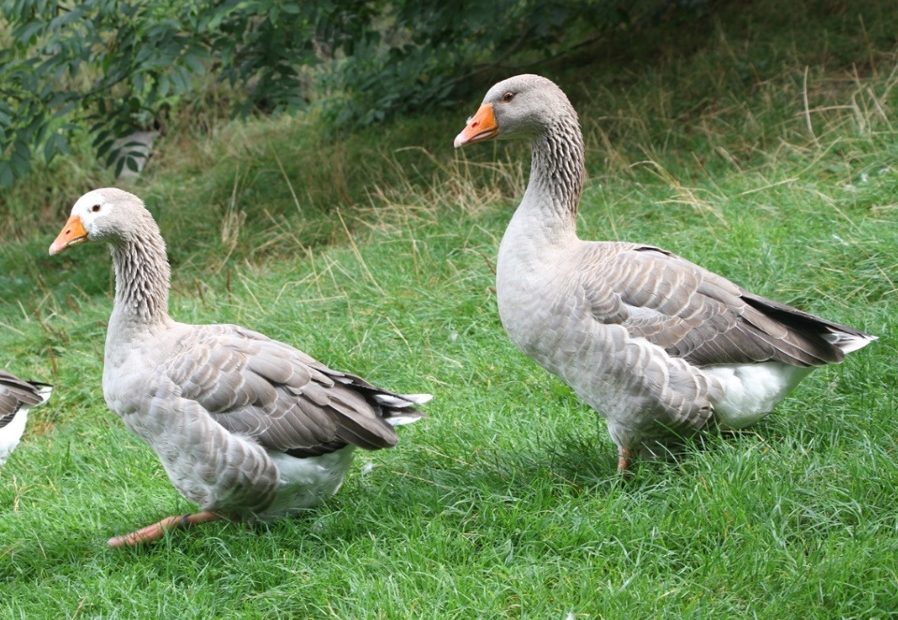  Pilgrim male x American Buff female F1-invidual. Carries also a recessive buff allele. Photo: Chris & Mike Ashton |
| T/G | ♂ -1 bp/C  Sd/sd+ |  | No photo available. We predict light plumage colour, but darker than in the genotype: G/G in *EDNRB2* and -1bp/C in *MLANA*. |
| T/G | ♂ -1 bp/-1 bp  Sd/Sd | autosexing white male, ’whiter’ overall than pure T/T males, heterozygosity often goes unnoticed, pink blood quills on the wing tips which show at 4-5 weeks | 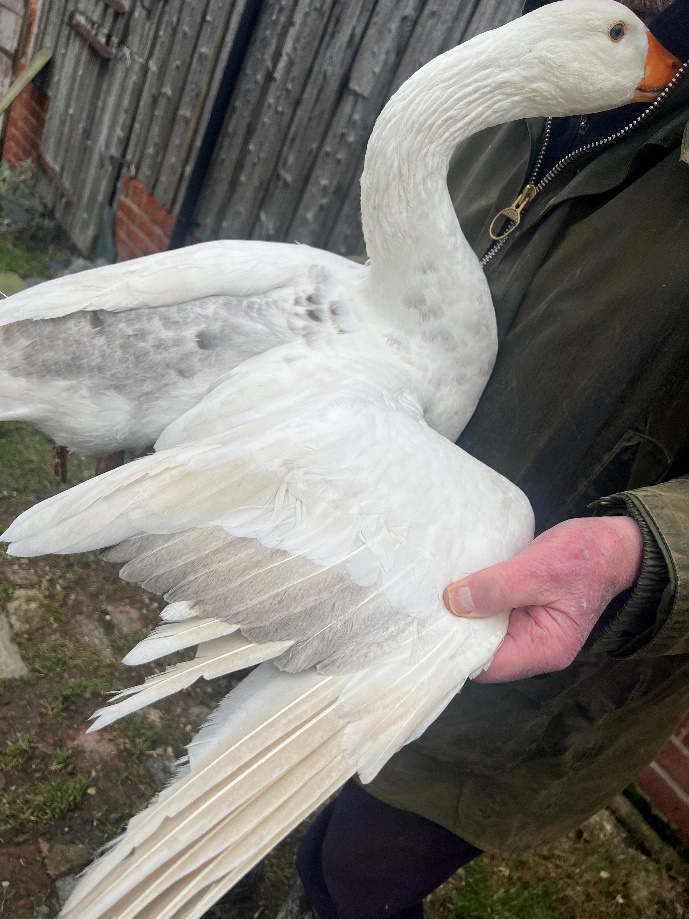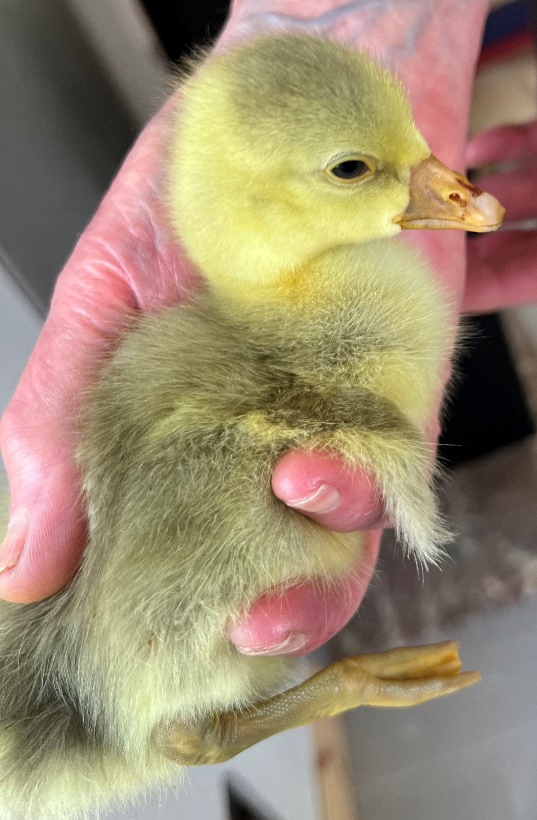  The wing tips of this gosling looked grey, instead of yellow, underbelly was yellow. He is mainly white, but the wing tips of flight feathers show some grey. Two outer flight feathers are white, which is an indication of spot carrier in Pilgrim breed.  Progeny of Shetland male x grey spot carrying female (Shetland male x Brecon Buff female). Sibling of geese in the next two photos. Photo: Chris & Mike Ashton  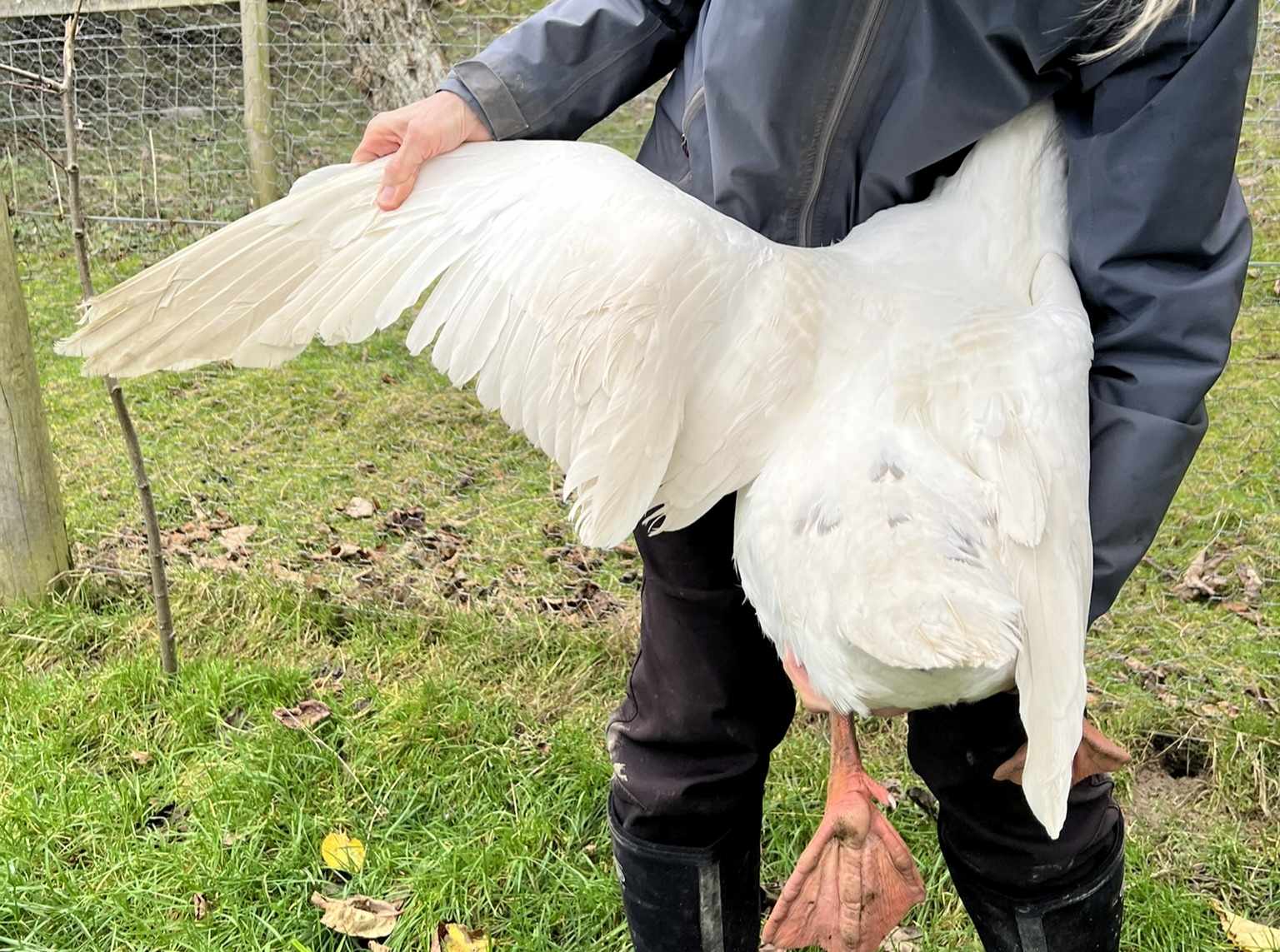  Pilgrim male x Czech female F1-individual. Photo: Chris & Mike Ashton |
| T/G | ♀ -1 bp  -/Sd | autosexing light grey female, females heterozygous for spotting show white chin spot/white flights/white breast patch (T/G is very variable in its expression) | 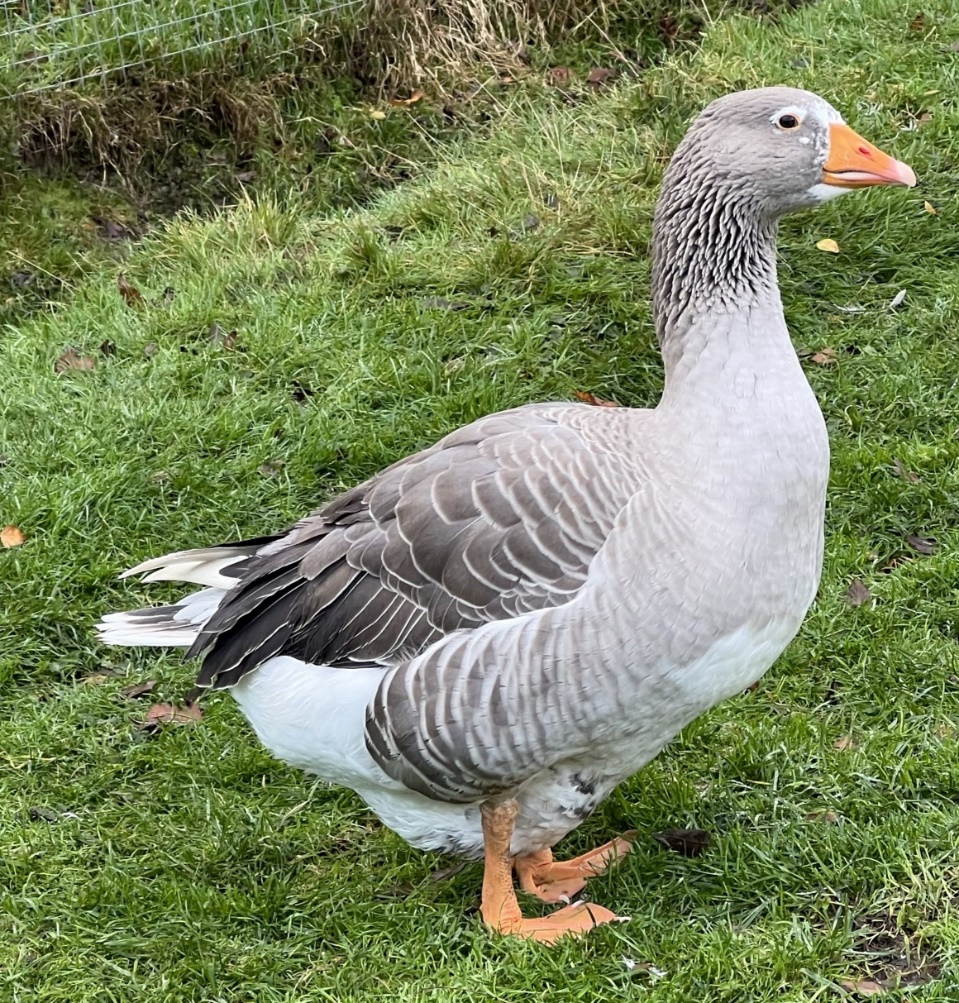  Shetland male x Brecon Buff female F1-individual. Photo: Chris & Mike Ashton  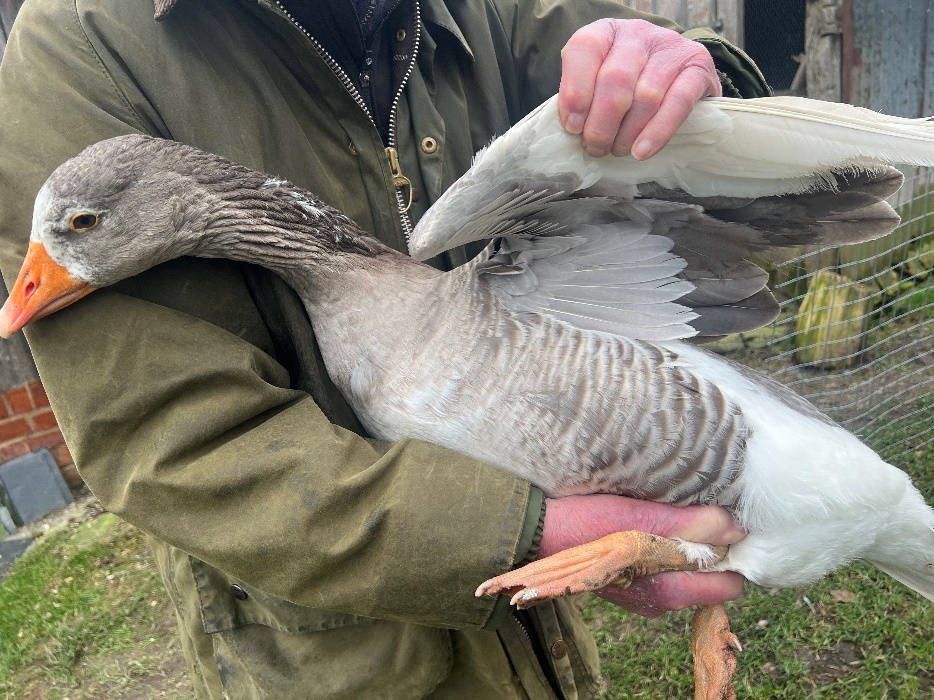  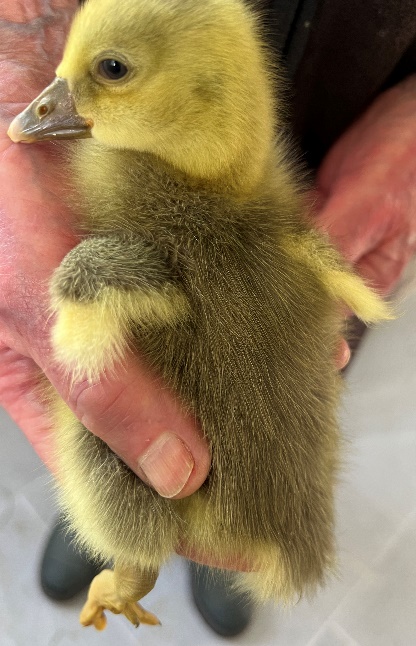  White spotting manifests in this gosling as yellow fluff in the wing tips and underbelly. In adult plumage she has a white breast patch, and several white fights on both wings.  Progeny of Shetland male x grey spot carrying female (Shetland male x Brecon Buff female). Sibling of geese in the upper and lower photos. Photo: Chris & Mike Ashton  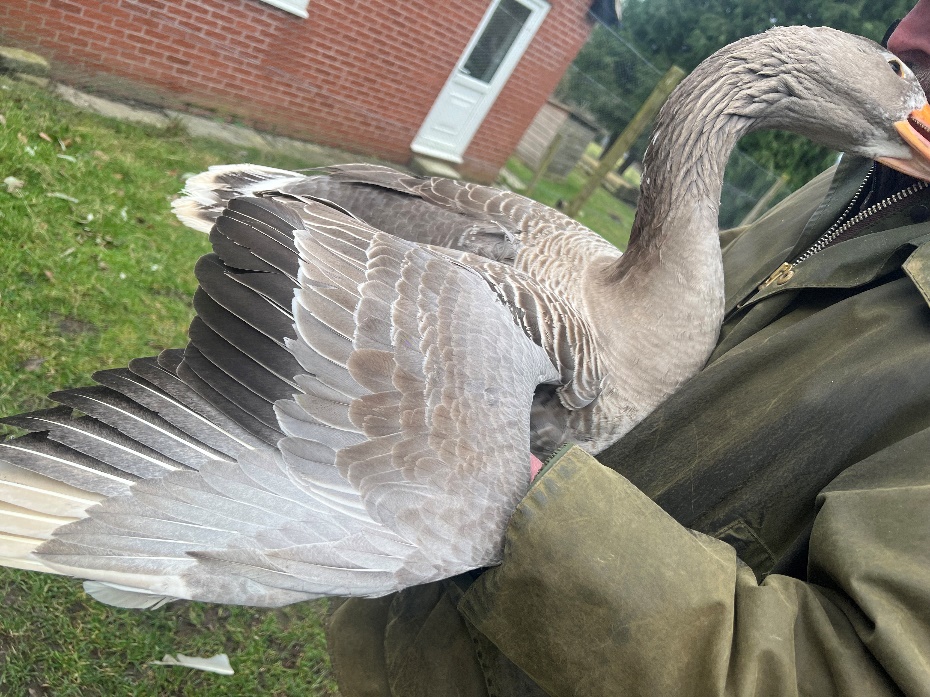  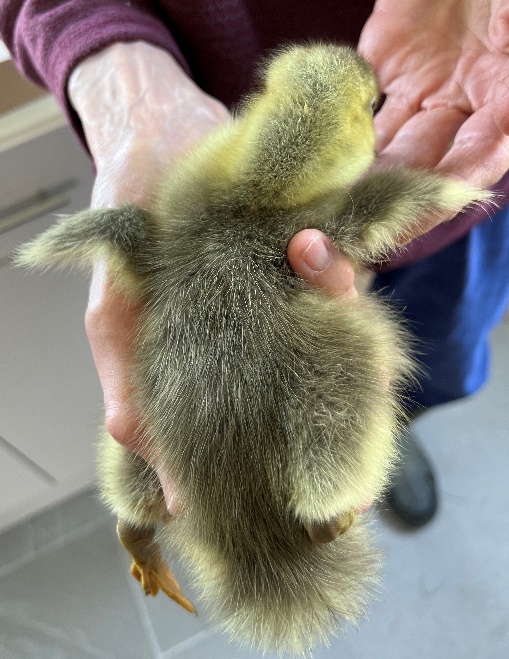  Gosling has less yellow compared to her sibling (in upper photo). In adult plumage she has four white flight feathers in her right wing, all left-wing primary feathers are nearly grey. Additionally, she has white chin spot.  Progeny of Shetland male x grey spot carrying female (Shetland male x Brecon Buff female). Sibling of geese in the upper two photos. Photo: Chris & Mike Ashton |
| G/G | ♂ C/C or ♀ C  sd+/sd+ or -/sd+ | saddleback (also called as pied/piebald/mottled (variable expression or another locus affects the amount of white spotting) | 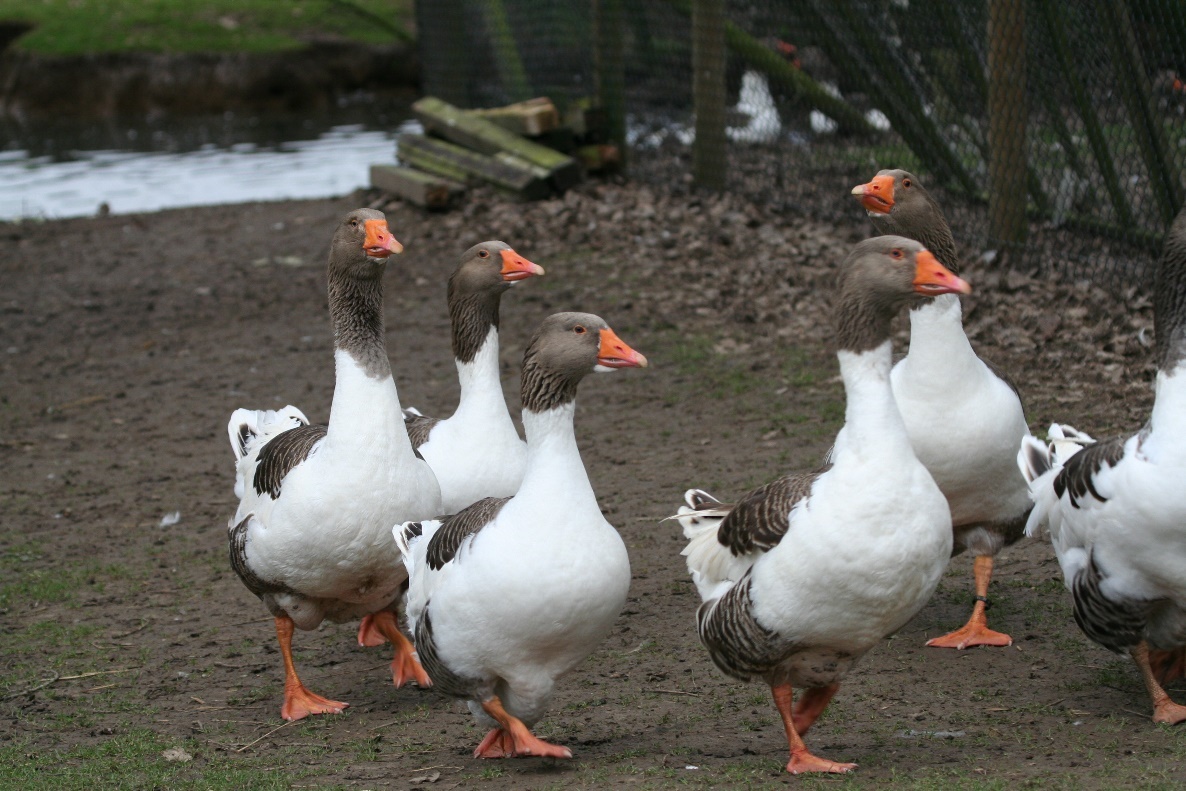  Pomeranian geese. Photo: Chris & Mike Ashton  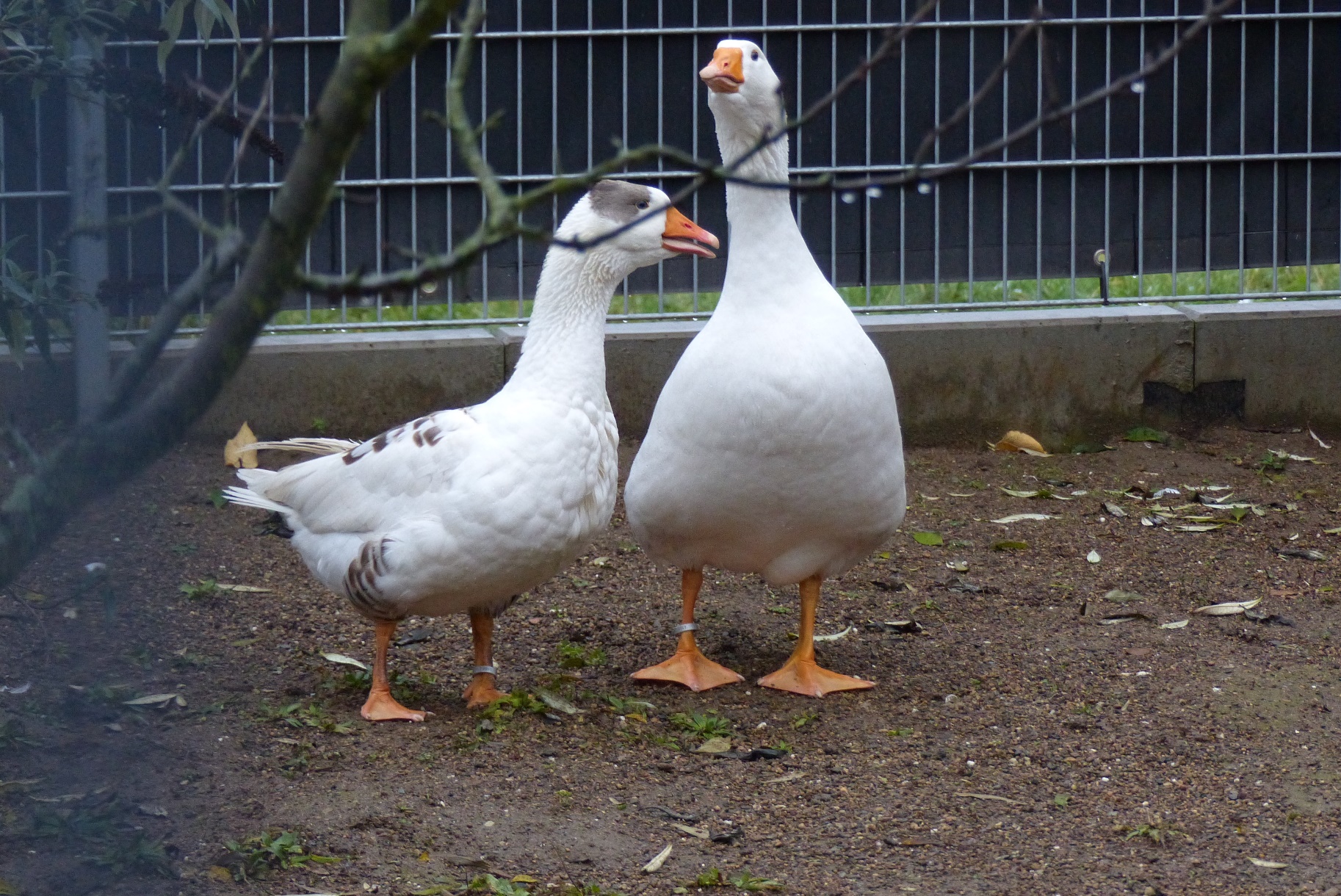  Leine goose female (sample no JS80 in this study). Note the large extent of white spotting, which could be caused by another locus. Photo: Maximilian Birkendorf, Zoo Neuwied  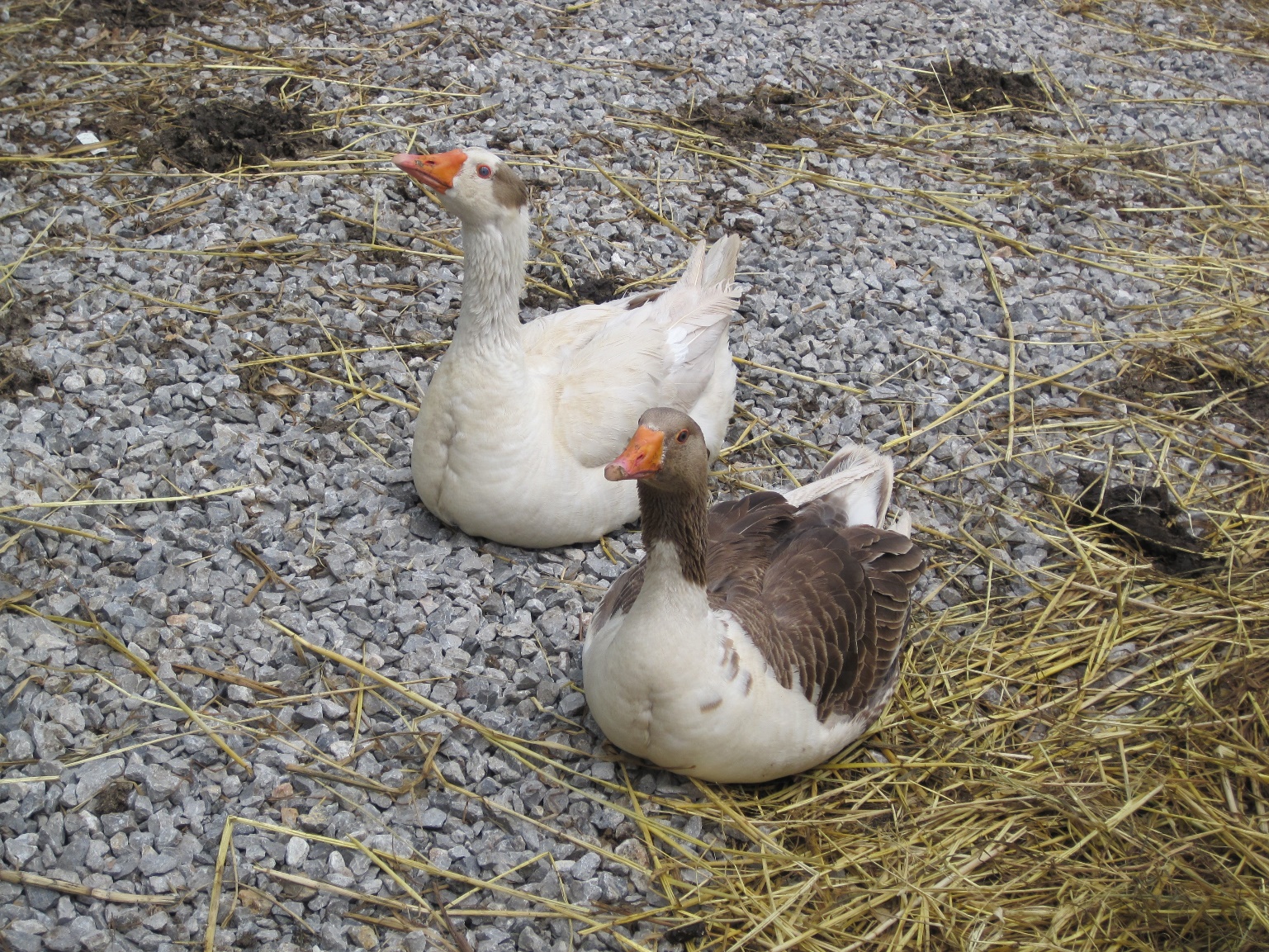  Non-breed Turkish geese (upper goose sample no TR2 in this study). Note a large extent of white spotting which could be caused by another locus. Photo: Marja E. Heikkinen  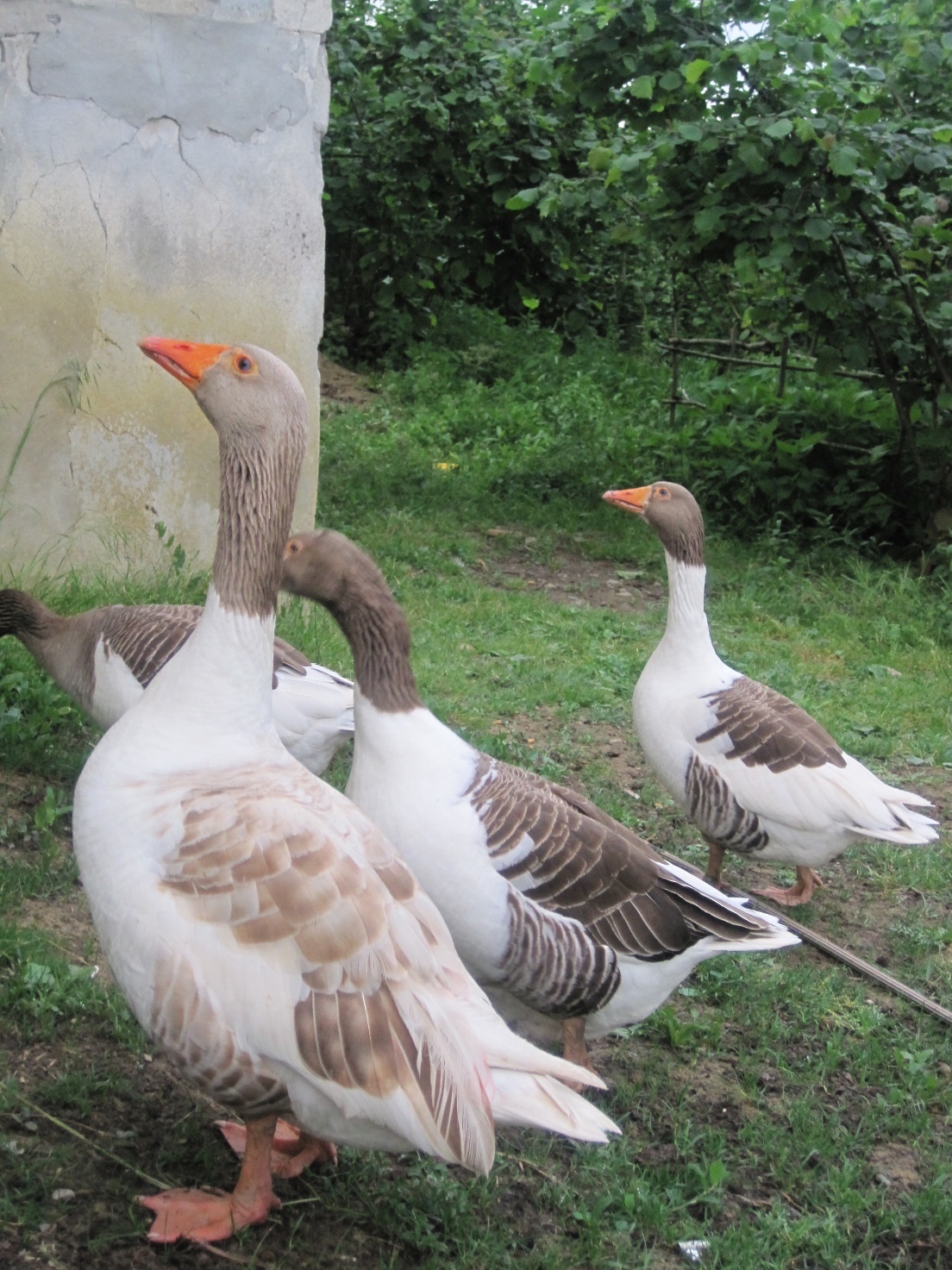  Non-breed Turkish geese with buff back colouration (goose in the foreground sample no TR5 in this study). Photo: Marja E. Heikkinen |
| G/G | ♂ -1 bp/C  Sd/sd+ | minimal grey spotting in thigh to saddleback pattern (variable expression or another locus affects the amount of white spotting) | 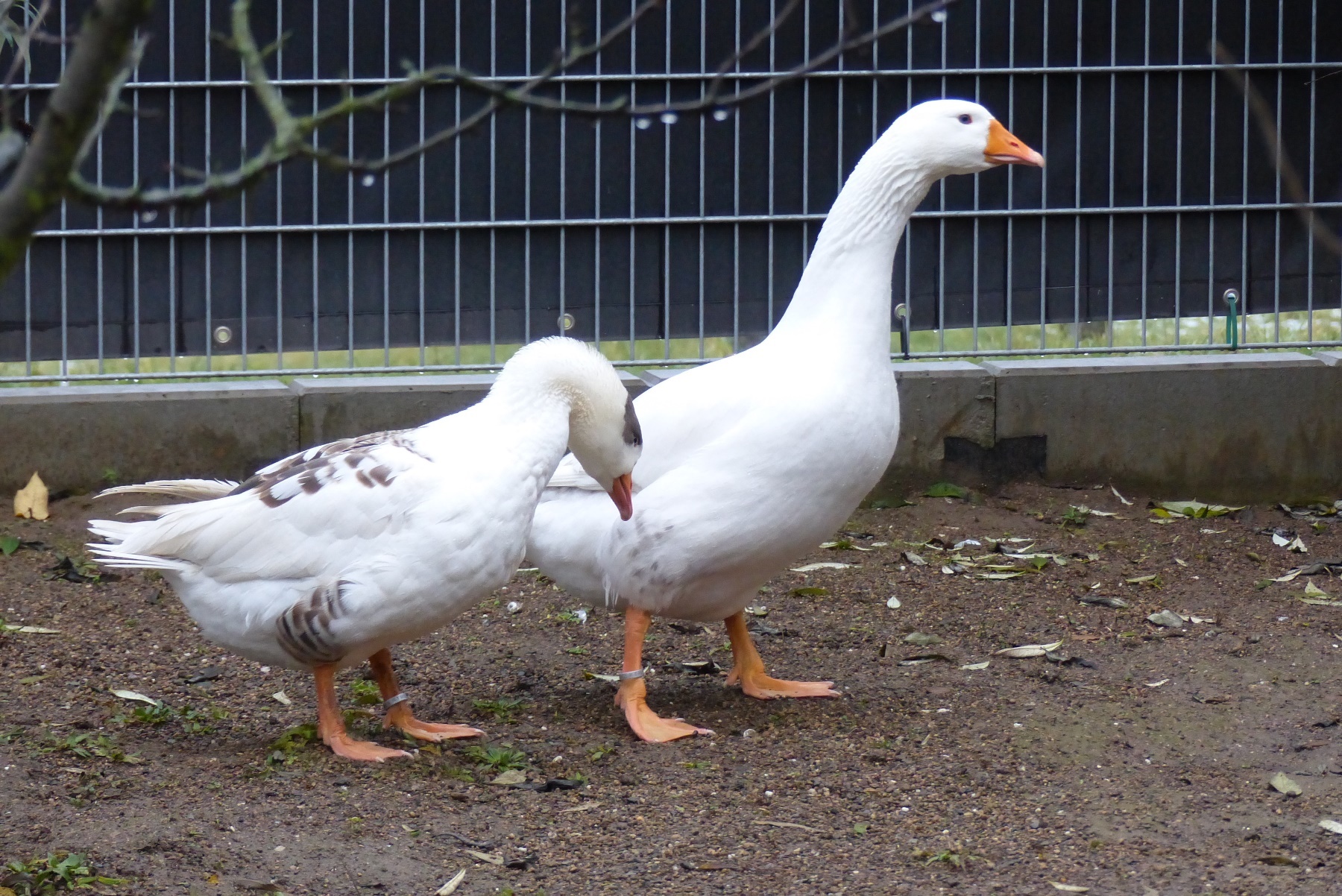  Leine goose male (sample no JS79 in this study). Photo: Maximilian Birkendorf, Zoo Neuwied |
| G/G | ♂ -1 bp/-1 bp or ♀-1 bp  Sd/Sd or -/Sd | white  OR autosexing almost white male and saddleback female, West of England, Shetland and Normandy breeds (probably have another locus which allows the spot to show in females, probably sex-linked) | 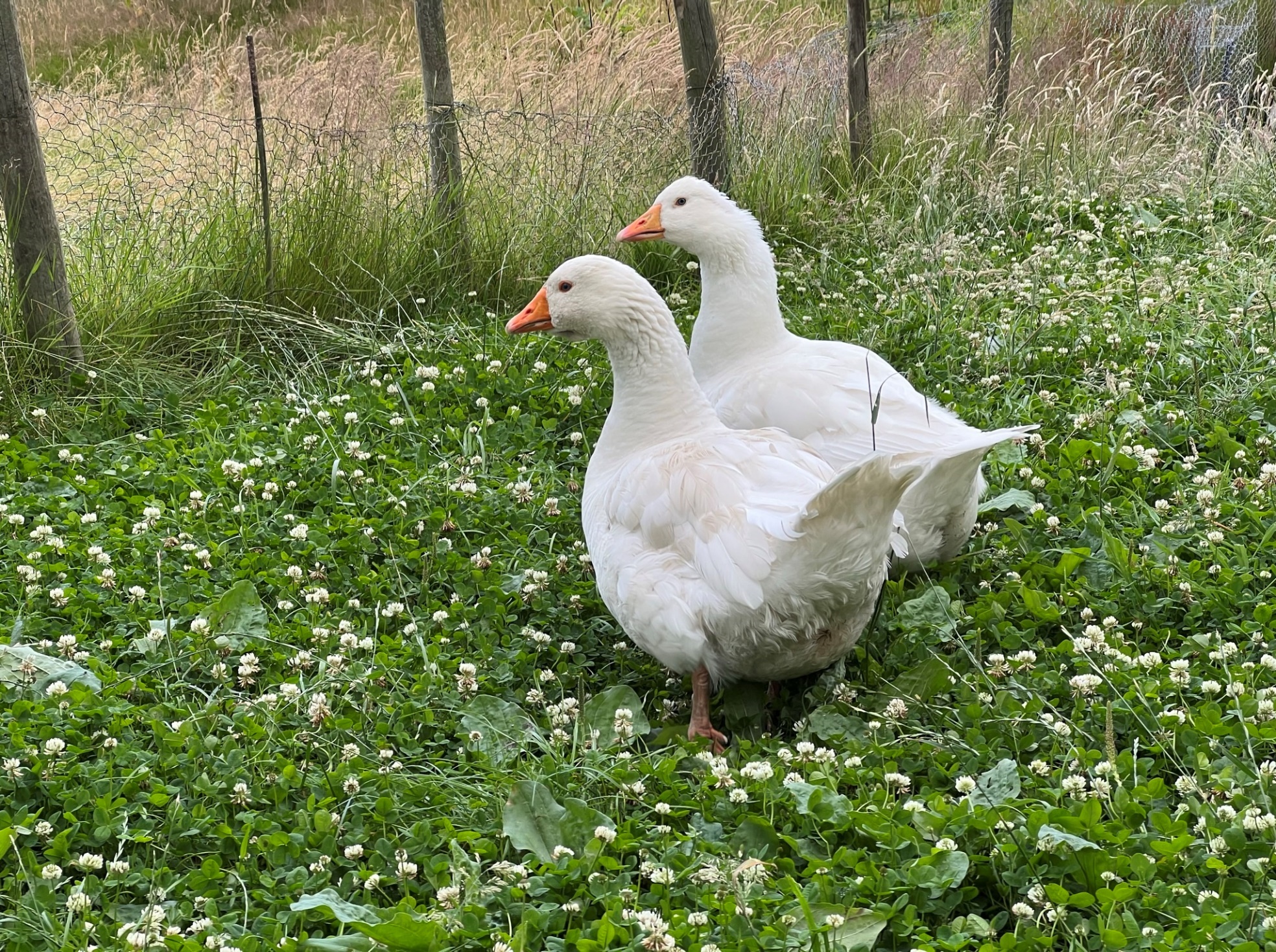  Czech geese. Photo: Chris & Mike Ashton  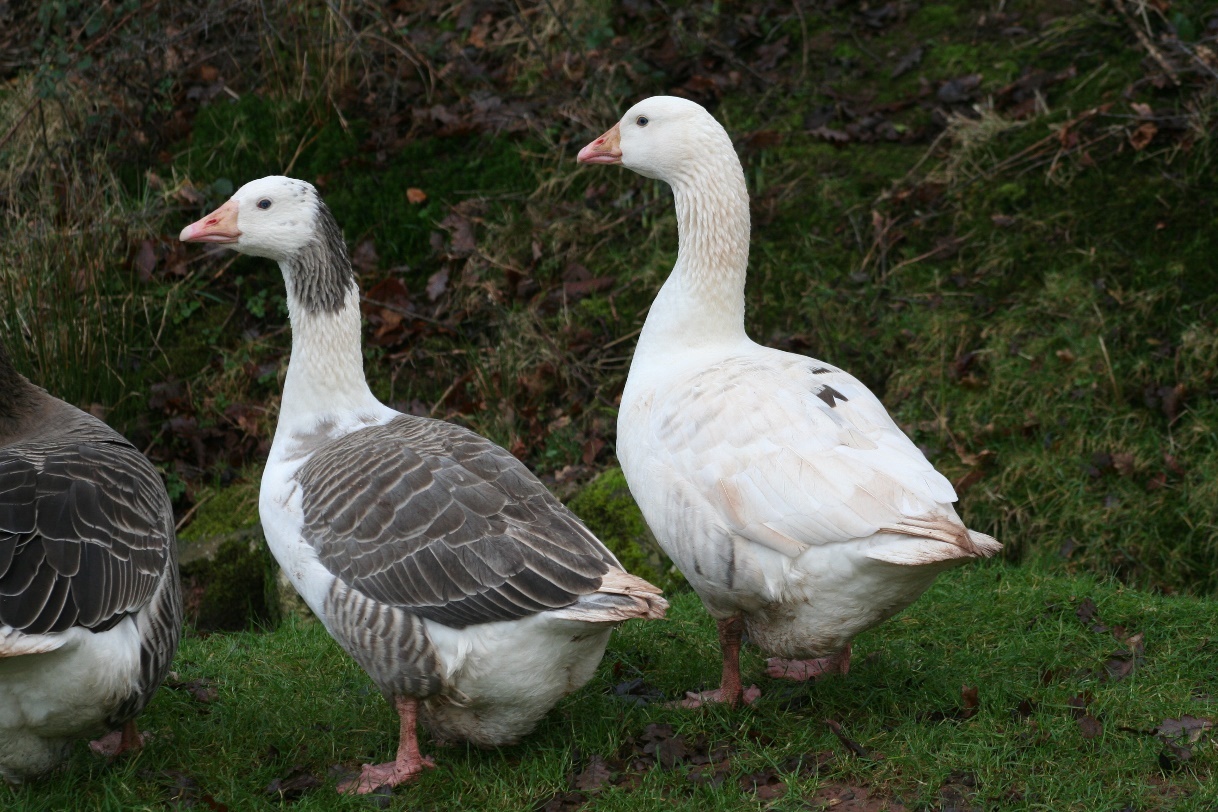  Shetland geese. Photo: Chris & Mike Ashton |

Genotypes in exon 3 of the gene *endothelin receptor B-like* (*EDNRB2*) known to affect colour polymorphism in domestic geese of swan goose origin (*Anser cygnoid*).

| **EDNRB2, position NW_013185915.1: g. 750,748–750,735 insertion** | **Phenotype** | **Photo** |
| --- | --- | --- |
| -/-  C/C | brown (aka grey) | 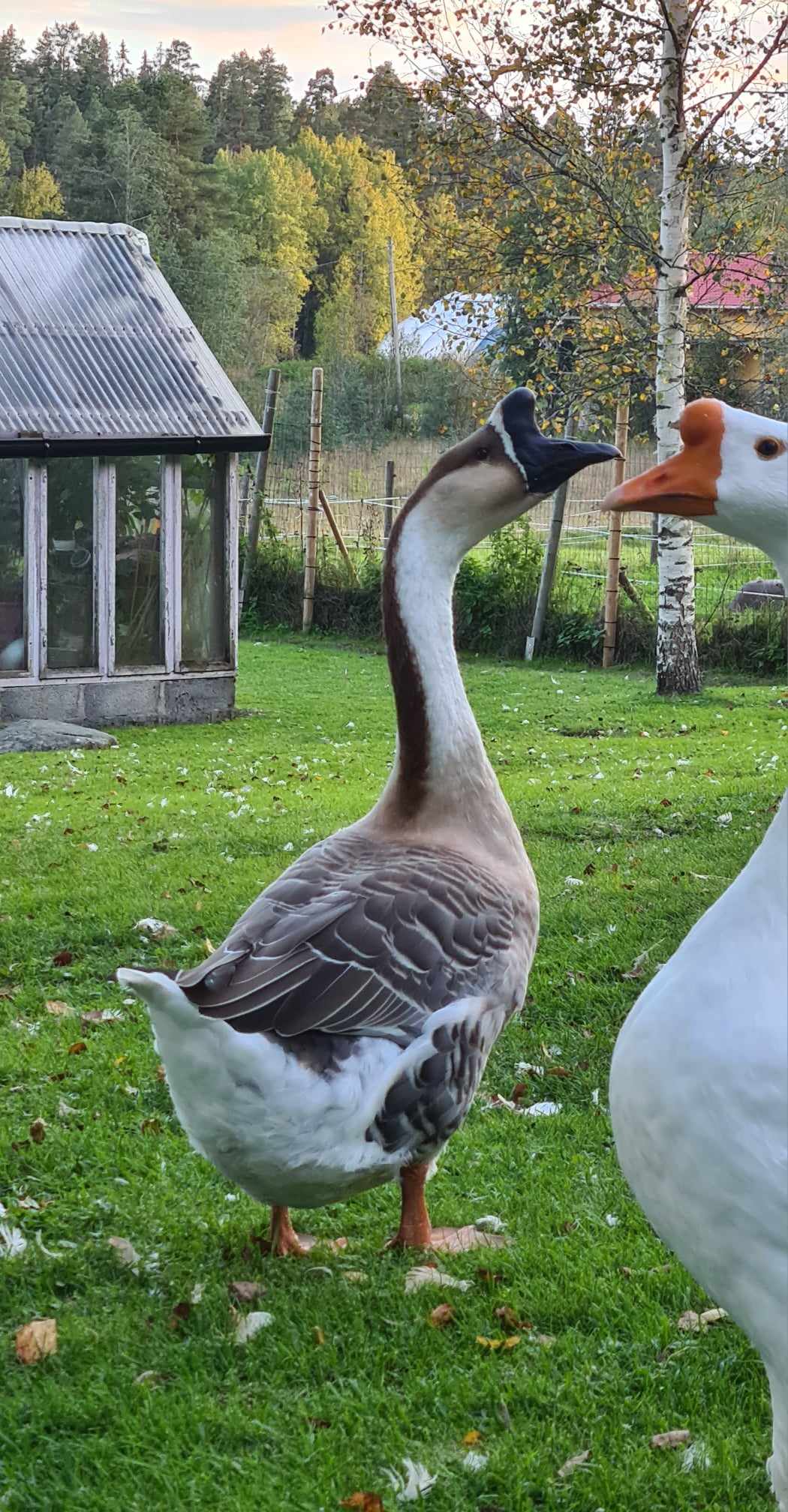  Chinese domestic goose (sample no JS70 in this study). Photo: Kirsi Mäki |
| -/14 bp insertion  C/c | brown (aka grey), with white breast pattern of variable size which can extend to the dorsal side and white primary flight feathers | 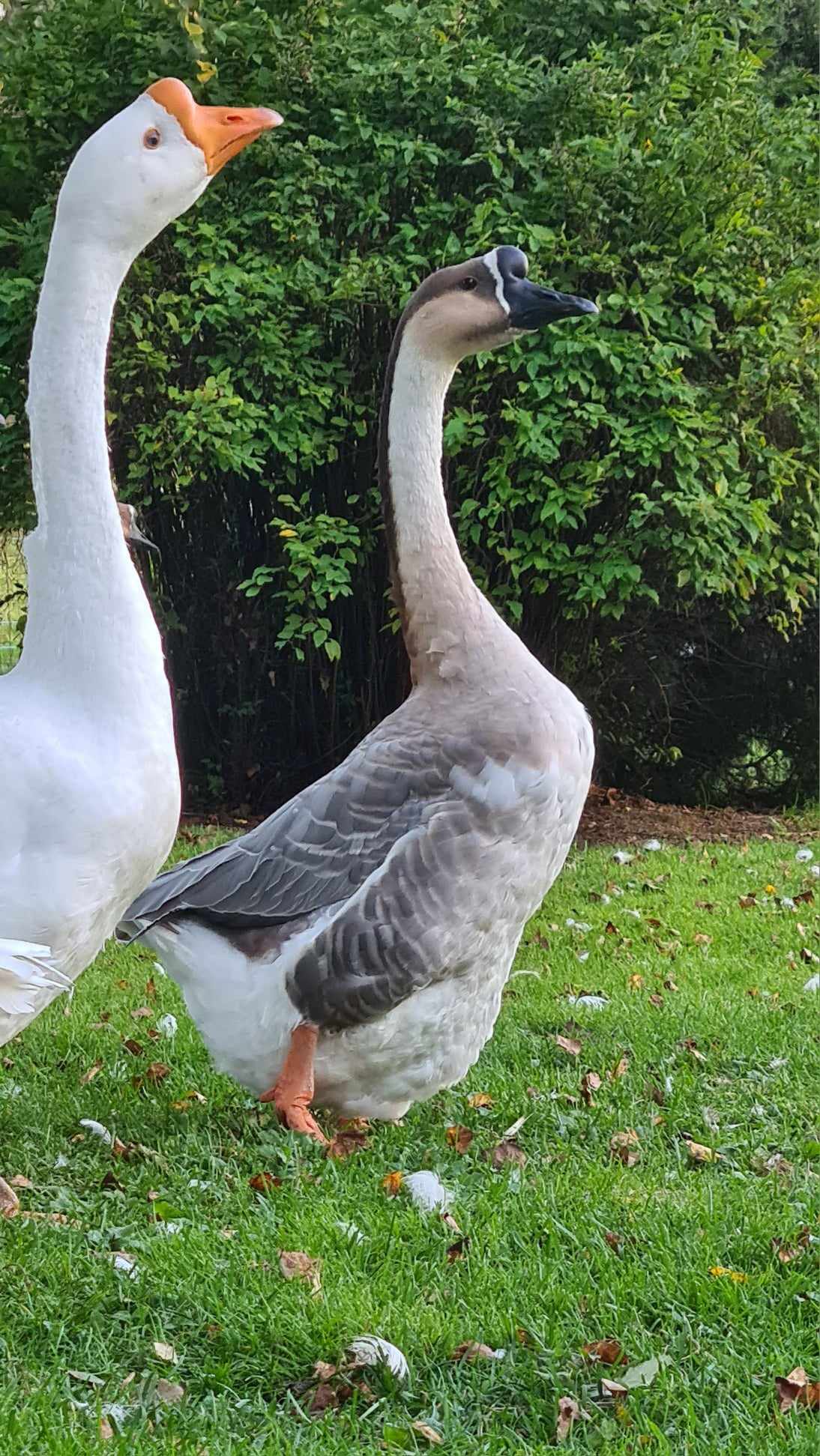  Chinese domestic goose, with also mutation for blue colour (sample no JS72 in this study). Photo: Kirsi Mäki |
| 14 bp insertion/14 bp insertion  c/c | white (small grey spots in the back, not visible when the wings are closed) | 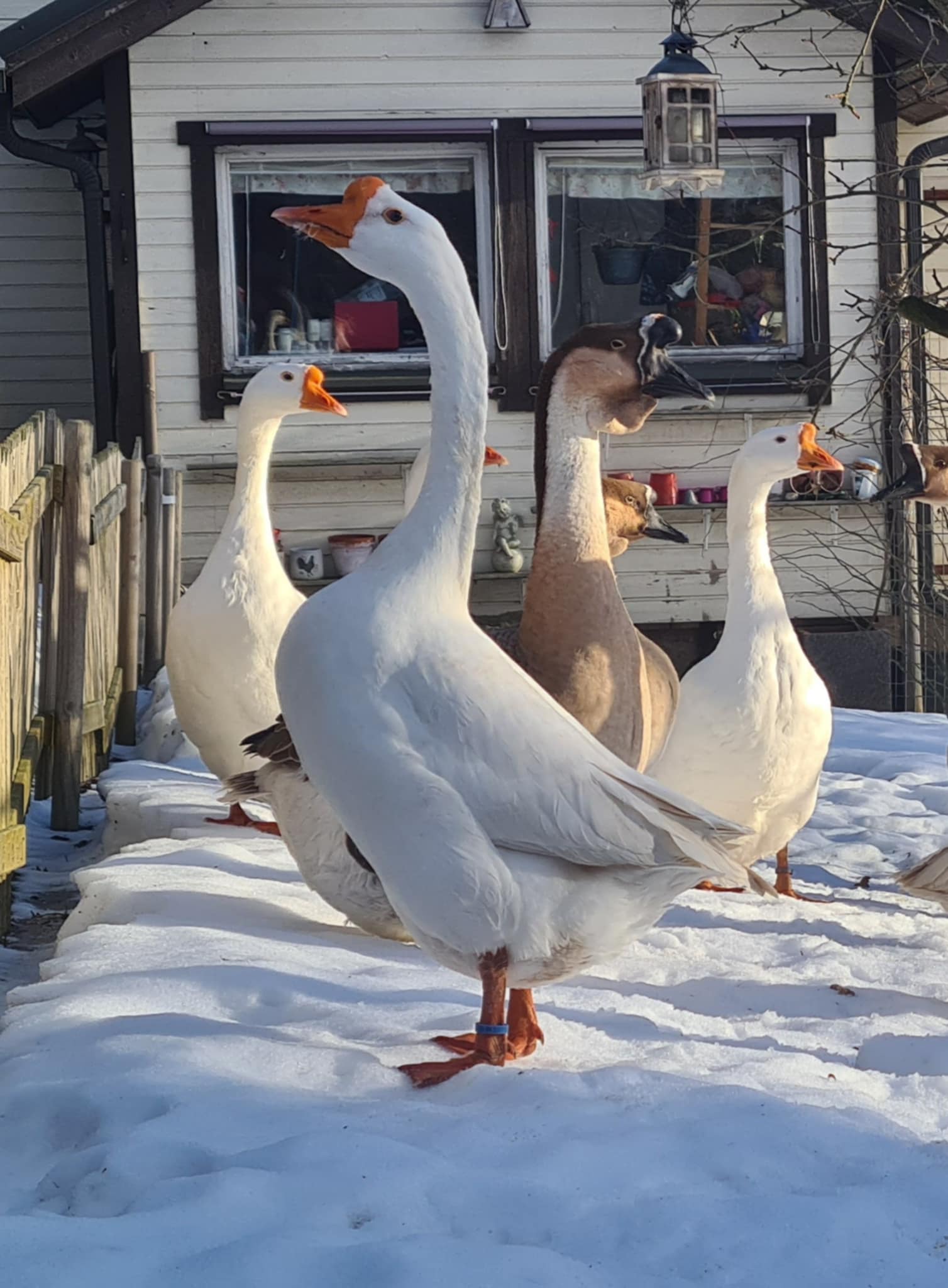  Chinese domestic goose. Photo: Kirsi Mäki |
